# Control of neurogenic competence in mammalian hypothalamic tanycytes

**DOI:** 10.1101/2020.11.02.359992

**Authors:** Sooyeon Yoo, Juhyun Kim, Pin Lyu, Thanh V. Hoang, Alex Ma, Vickie Trinh, Weina Dai, Lizhi Jiang, Patrick Leavey, Jae-Kyung Won, Sung-Hye Park, Jiang Qian, Solange P. Brown, Seth Blackshaw

## Abstract

Hypothalamic tanycytes, radial glial cells that share many features with neuronal progenitors, generate small numbers of neurons in the postnatal hypothalamus, but the identity of these neurons and the molecular mechanisms that control tanycyte-derived neurogenesis are unknown. In this study, we demonstrate that tanycyte-specific disruption of the NFI family of transcription factors (*Nfia/b/x*) robustly stimulates tanycyte proliferation and tanycyte-derived neurogenesis. Single-cell RNA- and ATAC-Seq analysis reveals that NFI factors repress Shh and Wnt signaling in tanycytes, and small molecule modulation of these pathways blocks proliferation and tanycyte-derived neurogenesis in *Nfia/b/x*-deficient mice. We show that *Nfia/b/x*-deficient tanycytes give rise to multiple mediobasal hypothalamic neuronal subtypes that can mature, integrate into hypothalamic synaptic circuitry, and selectively respond to changes in internal states. These findings identify molecular mechanisms that control tanycyte-derived neurogenesis, suggesting a new therapeutic approach to selectively remodel the hypothalamic neural circuitry that controls homeostatic physiological processes.

## Introduction

Hypothalamic tanycytes are radial glial cells that line the ventricular walls of the mediobasal third ventricle (*1*, *2*). Tanycytes are subdivided into alpha1, alpha2, beta1 and beta2 subtypes based on dorso-ventral position and marker gene expression, and closely resemble neural progenitors in morphology and gene expression profile. Tanycytes have been reported to generate small numbers of neurons and glia in the postnatal period, although at much lower levels than in more extensively characterized sites of ongoing neurogenesis, such as the subventricular zone of the lateral ventricles, or the subgranular zone of the dentate gyrus (*3*–*6*). While tanycyte-derived newborn neurons may play a role in regulating a range of behaviors (*3*, *7*, *8*), levels of postnatal tanycyte-derived neurogenesis are low and virtually undetectable in adulthood. As a result, little is known about the molecular identity or connectivity of tanycyte-derived neurons (*6*, *9*). A better understanding of the gene regulatory networks that control neurogenic competence in hypothalamic tanycytes would both give insight into the function of tanycyte-derived neurons and potentially identify new therapeutic approaches for modulation and repair of hypothalamic neural circuitry.

Studying retinal Müller glia, which closely resemble hypothalamic tanycytes in both morphology and gene expression, provides valuable insight into the neurogenic potential of tanycytes (*1*, *3*, *9*, *10*). Zebrafish Müller glia function as quiescent neural stem cells, and are able to regenerate every major retinal cell type following injury (*11*). While mammalian Müller glia effectively lack neurogenic competence, in posthatch chick they retain a limited neurogenic competence that resembles that of mammalian tanycytes (*12*, *13*). Recent studies in retina have identified the NFI family of transcription factors *Nfia/b/x* as being essential negative regulators of neurogenesis in both late-stage progenitor cells and in mature mammalian Müller glia (*14*, *15*). Moreover, like in retina, NFI factors are expressed in late-stage hypothalamic neural progenitors (*16*), and *Nfia* is necessary for hypothalamic glia specification (*17*). These findings raise the possibility that NFI factors may actively repress proliferation and

We hypothesize here that suppression of NFI factor activity in tanycytes may enhance their proliferative and neurogenic capacity. To address this possibility, we selectively disrupted *Nfia/b/x* function in hypothalamic tanycytes of both juvenile and adult mice. We observed that early loss of function of NFI activity in hypothalamic tanycytes led to a robust induction of proliferation and neurogenesis, while *Nfia/b/x* disruption in adults led to lower levels of tanycyte-derived proliferation than are seen following neonatal loss of function. NFI loss of function activated both Shh and Wnt signaling in tanycytes, and this in turn stimulated proliferation and neurogenesis. These tanycyte-derived neurons survive, mature, migrate radially away from the ventricular zone, express molecular markers of diverse hypothalamic neuronal subtypes, and functionally integrate into hypothalamic synaptic circuitry. These findings demonstrate that hypothalamic tanycytes possess a latent neurogenic competence that is actively suppressed by NFI family transcription factors, which can be modulated to induce generation of multiple hypothalamic neuronal subtypes.

## Results

### *Nfia/b/x* loss of function induces proliferation and neurogenesis in neonatal mice

To determine whether hypothalamic tanycytes express *Nfia/b/x*, we first analyzed bulk RNA-Seq data obtained from FACS-isolated GFP-positive cells from the mediobasal hypothalamus of adult Rax-GFP transgenic mice, where GFP is selectively expressed in tanycytes (*18*, *19*). We observe that GFP-positive tanycytes, indicated by *Rax* expression, show highly enriched expression for *Nfia, Nfib*, and *Nfix* relative to the GFP-negative, neuronally-enriched fraction of mediobasal hypothalamic cells (Fig. 1A). Immunohistochemical analysis in adult mice reveals that Nfia/b/x proteins are highly expressed in the tanycytic layer and in other hypothalamic glial cells in adult mice (Fig. 1B) (*17*). We next generated *Rax-CreER;Nfia^lox/lox^;Nfib^lox/lox^;Nfix^lox/lox^;CAG-lsl-Sun1-GFP* mice (TKO mice hereafter), which allow inducible, tanycyte-specific disruption of *Nfia/b/x* function while simultaneously tracking the fate of tanycyte-derived cells using Cre-dependent Sun1-GFP expression (Fig. 1C) (*14*, *18*, *20*).

**Figure 1:**
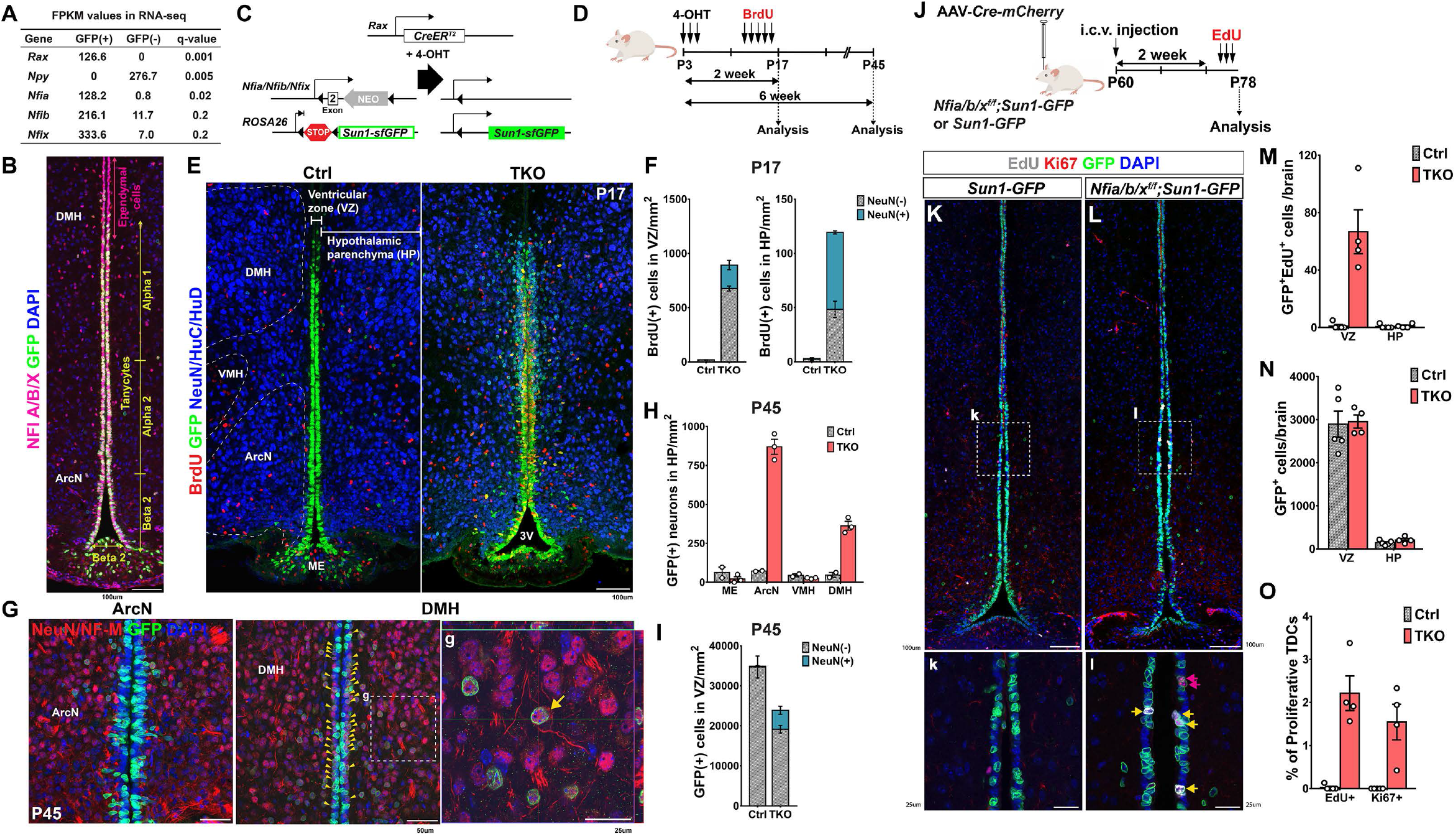
*Nfia/b/x* suppress proliferation and neurogenesis in tanycytes of neonatal mice. A. Expression of *Nfia/b/x* in GFP+ tanycytes isolated from Rax:GFP mice (*19*) compared to the GFP-negative cells in adult hypothalamus. The tanycyte-specific marker *Rax* and the neuronal marker *Npy* are enriched in GFP+ and GFP-cells, respectively. B. Distribution of Nfia/b/x protein in Rax-GFP+ tanycytes. C. Schematic of mouse lines used in this study. D. Schematic of a genetic approach for simultaneous tanycyte-specific disruption of *Nfia/b/x* and reporter gene labeling of tanycytes and tanycyte-derived cells using tamoxifen-dependent activation of CreER. E. Induction of proliferation and neurogenesis in NFI-deficient tanycytes by P17. F. Quantification of proliferation and neurogenesis in the ventricular zone (VZ) and hypothalamic parenchyma (HP) at P17 (n=3-5 mice). G. In NFI TKO mice by P45, NeuN/neurofilament M-positive tanycyte-derived neurons migrate into the parenchyma of the arcuate nucleus (ArcN) and dorsomedial hypothalamus (DMH), with a small number of neurons remaining in the subventricular zone (yellow arrowheads). Enlarged image of parenchymal tanycyte-derived neurons in g with the orthogonal view showing co-staining within the cell. H. Substantially increased numbers of NeuN+/GFP+ tanycyte-derived neurons are observed in NFI TKO mice in ArcN and DMH relative to wildtype controls, but comparable numbers of neurons are observed in median eminence (ME) and ventromedial hypothalamus (VMH) (n=2-3 mice). I. The number of GFP+ tanycytes is reduced in NFI-deficient mice at P45, and ectopic neurons are seen in the VZ (n=2-3 mice). J. Schematic for i.c.v. delivery of AAV-Cre and analysis of *Nfia/b/x* loss of function in P78 mice. K. AAV-Cre induces Sun1-GFP expression in tanycytes in *CAG-lsl-Sun1-GFP* control mice at P78 (k inset shows alpha tanycytes). L. AAV-Cre induces proliferation in alpha tanycytes (shown in inset l) of *Nfia^lox/lox^;Nfib^lox/lox^;Nfix^lox/lox^;CAG-lsl-Sun1-GFP* mice. M. Quantification of GFP+/EdU+ cells in VZ and HP in NFI-deficient mice (n=4-5 mice). N. Number of GFP+ cells in VZ and HP in control and NFI-deficient mice. O. Percentage of GFP+ VZ cells in the alpha tanycyte region labeled by Ki67 and EdU in NFI-deficient mice (n=4-5 mice). Scale bars: B,E,K,L=100 μm; G=50 μm; g,k,l=25 μm.

We induced Cre activity using daily intraperitoneal (i.p.) injections of 4-hydroxytamoxifen (4-OHT) between postnatal day 3 and day 5 (P3 and P5) (Fig. 1D). At this point, neurogenesis in the mediobasal hypothalamus is low under baseline conditions (*3*, *9*). Following 4-OHT treatment, we observe that NFIA/B/X immunoreactivity is first reduced in the tanycyte layer beginning at P6 following 4-OHT injections between P3 and P5, initially in more ventral regions where Rax expression is strongest (Fig. S1A). NFIA/B/X immunoreactivity is largely undetectable by P10, and Cre-dependent GFP expression was correspondingly induced (Fig. S1A). BrdU incorporation and Ki67 labeling was seen beginning at P6 in dorsally-located alpha tanycytes, with labeling spreading to beta1 tanycytes of the arcuate nucleus by P8, and beta2 tanycytes of the median eminence by P10 (Fig. S1A, S1B). At P12, Ki67 labeling was observed in the tanycyte layer immediately adjacent to the third ventricle lumen, and a small subset of GFP-positive cells in the tanycyte layer closest to the hypothalamic parenchyma began to express neuronal markers (Fig. S1C).

By P17, Nfia/b/x expression was completely lost in the tanycytes, although it was preserved in Rax-negative ependymal cells where GFP expression is not induced (Fig. S1D), and tanycyte-derived GFP-positive neuronal precursors in the ventricular zone were actively amplified and had begun to migrate outward into the hypothalamic parenchyma in *Nfia/b/x*-deficient mice (Fig. 1E,F, S1D). In contrast, few migrating GFP-positive cells were observed in *Rax-CreER;CAG-lsl-Sun1-GFP* controls (Fig. 1E,F). At P45, a substantial increase in GFP-positive cells expressing mature neuronal markers was observed in the parenchyma of the arcuate (ArcN) and dorsomedial (DMH) hypothalamic nuclei (Fig. 1G,H), while limited numbers of GFP-positive cells expressing neuronal markers remained in the subventricular region in *Nfia/b/x*-deficient mice (Fig. 1G, yellow arrowheads, and 1I).

### Loss of function of *Nfia/b/x* induces tanycyte proliferation in older mice

Our previous studies showed that loss of *Nfia/b/x* in late-stage retinal progenitors robustly induces proliferation and neurogenesis under baseline conditions, while in adult Müller glia, it induces limited levels of proliferation and neurogenesis, but only following neuronal injury (*14*, *15*). This suggests that neurogenic competence in mature murine tanycytes could be lower than that seen in neonates. To test this, we conducted 4-OHT treatment at older ages. While we still observed a robust induction of proliferation and neurogenesis following treatment at P7, this was less effective at P10. Only very low levels were observed at P12, not significantly different from control mice (Fig. S2A,B,C). However, this low level of proliferation reflected a substantially reduced efficiency of 4-OHT-dependent disruption of *Nfia/b/x* as confirmed by the largely intact pattern of immunoreactivity for NFIA/B/X in TKO mice (Fig. S2D).

To improve the efficiency of *Nfia/b/x* deletion, and to study the effects of NFI loss of function in adult animals, we applied viral-mediated Cre delivery. Intracerebroventricular (i.c.v.) injection of AAV1-Cre-mCherry into both *Nfia^lox/lox^;Nfib^lox/lox^;Nfix^lox/lox^;CAG-lsl-Sun1-GFP* and *CAG-lsl-Sun1-GFP* control mice at P60 resulted in robust mCherry expression in ventricular hypothalamic cells within two weeks, along with Cre-dependent induction of Sun1-GFP expression in both control and *Nfia/b/x* floxed conditional mice (Fig. S3). We observed efficient loss of NFI expression in GFP-positive tanycytes by P74 and co-immunolabeling with BrdU, delivered continuously by osmotic mini-pump during the two weeks (Fig. S3C). To determine the specific induction of proliferation initiated from tanycytes, we administered EdU using a once daily i.p. injection between P76-P78, and analyzed mice at P79. We observed selective EdU incorporation into alpha tanycytes adjacent to the dorsal part of ArcN (Fig. 1L, l, M). Much lower levels of EdU incorporation were observed in ventrally located beta tanycytes, while no EdU labeling was observed in controls (Fig.1K, k). Although we observed a few GFP-positive cells in the mediobasal hypothalamic parenchyma, these cells were not labeled with EdU, and there was no difference in their number between control and TKO animals (Fig. 1N). We confirmed the specific and local induction of tanycyte proliferation with the increased endogenous expression of Ki67, a proliferation marker, only in *Nfia/b/x*-deficient TKO mice (Fig. 1O, S3C).

### Single cell RNA-Seq and ATAC-Seq analysis identify gene regulatory networks controlling neurogenesis in tanycytes

To obtain a comprehensive picture of the cellular and molecular changes that occur during proliferation and neurogenesis, we conducted scRNA-Seq analysis of FACS-isolated GFP-positive tanycytes and tanycyte-derived cells from both control and *Nfia/b/x*-deficient mice. To do this, we induced Cre activity in tanycytes between P3 and P5 and harvested GFP-positive cells at P8, P17 and P45 timepoints in TKO mice. We next profiled a total of >60,000 cells using the Chromium platform (10XGenomics), generated separate UMAP plots for controls, *Nfia/b/x*-deficient tanycytes, and then aggregated data obtained from all samples (Fig. 2A, S4). In control mice, we could readily distinguish tanycyte subtypes (alpha1, alpha2, beta1, beta2), based on previously characterized molecular markers (*21*–*23*), but also observed that tanycytes give rise to a range of other hypothalamic cell types (Fig. 2B). In controls, at P8, we observe a small fraction of proliferative tanycytes, from which arise differentiation trajectories that give rise to astrocytes and ependymal cells, as well as small numbers of oligodendrocyte progenitor cells and neurons (Fig. 2C,D, Fig. S4, Table ST2). At P17 and P45, however, very few proliferating tanycytes are observed, and evidence for ongoing generation of neurons and glia is lacking (Fig. 2C,D, Fig. S4). In TKO mice, in contrast, we observe that a significantly larger fraction of cells are proliferating tanycytes at all ages, along with clear evidence for ongoing neurogenesis (Fig. 2C, Fig. S4). Furthermore, a substantial reduction in the relative fraction of non-neuronal cells -- including astrocytes, oligodendrocyte progenitor cells, and ependymal cells -- is observed, in line with previous studies reporting an essential role for NFI family genes in gliogenesis in other CNS regions (*14*, *24*–*26*). While controls show a higher fraction of tanycyte-derived astrocytes, a much higher fraction of GFP-positive cells are neurons in TKO mice (Fig. 2D, Table ST2). Furthermore, the density and relative fraction of cells expressing alpha2 tanycyte markers is increased in TKO mice, which is consistent with previously published neurosphere and cell lineage analysis (*4*), demonstrating a higher neurogenic competence for alpha2 tanycytes (Fig. 2C bottom, 2D, Table ST2) (*4*).

**Figure 2:**
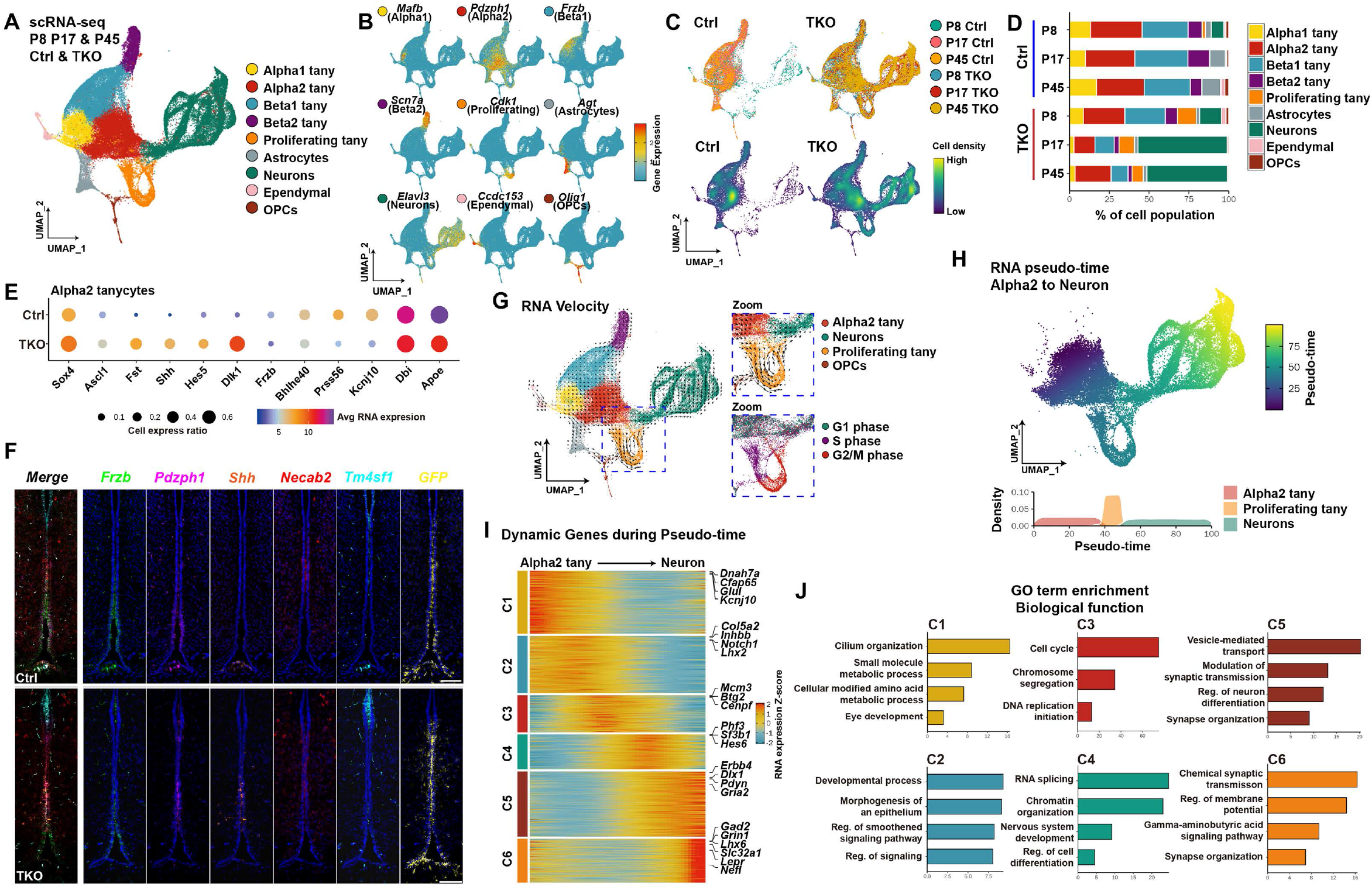
Single-cell RNA-Seq analysis of control and NFI-deficient tanycytes. A. Aggregate UMAP plot of scRNA-Seq data from control and NFI-deficient GFP+ tanycytes and tanycyte-derived cells isolated at P8, P17 and P45. Cell types are indicated by color shading. B. Distribution of cell type-specific marker expression on aggregate UMAP plot. C. Distribution of cells by age and genotype on aggregate UMAP plot. D. Percentage of each cell type by age and genotype. E. Dot plot showing genes differentially expressed in *Nfia/b/x*-deficient alpha2 tanycytes. F. Hiplex analysis for tanycyte subtype-specific markers and enhanced *Shh* expression in a subset of *Pdzph1+* alpha2 tanycytes. *Necab1+* alpha1 tanycytes and *Tm4sf1+* ependymal cells are shown for reference. G. RNA velocity analysis indicating differentiation trajectories in tanycytes and tanycyte-derived cells. Insets highlight proliferating tanycytes and tanycyte-derived neurons. H. Pseudotime analysis of differential gene expression in alpha2 tanycytes, proliferating tanycytes and tanycyte-derived neurons. I. Heatmap showing differentially expressed genes over the course of tanycyte-derived neurogenesis. J. Gene Ontology (GO) analysis of differentially expressed genes in I, with enrichment shown at −log10 P-value. Scale bars: F=100 μm. tany= tanycytes.

To identify critical regulators of proliferative and neurogenic competence in tanycytes, we performed a differential gene expression analysis between control and TKO mice in each tanycyte subtype. This analysis uncovered differential expression of multiple extrinsic and intrinsic regulators of these processes, particularly in alpha2 tanycytes (Fig. S5, Table ST4, Fig. 2E). Control tanycytes also selectively expressed many genes expressed by both mature, quiescent tanycytes and retinal Müller glia, whose expression is downregulated following cell-specific deletion of *Nfia/b/x (15*). These include genes that are highly and selectively expressed in mature alpha2 tanycytes, such as *Apoe* and *Kcnj10*, the Notch pathway target *Hes1*, the Wnt inhibitor *Frzb*, and the transcription factors *Klf6* and *Bhlh40. Nfia/b/x*-deficient tanycytes, in contrast, upregulated *Shh*, the Notch inhibitor *Dlk1*, the BMP inhibitor *Fst*, the neurogenic factors *Ascl1* and Sox4, and the Notch pathway target *Hes5* (Fig. 2E). Alpha1 and beta1 tanycytes also showed reduced expression of *Tgfb2* and *Fgf18* (Fig. S5), which were previously shown to be strongly expressed in these cells (*23*, *27*). To validate these results, we conducted multiplexed smfISH (HiPlex, ACD Bio-Techne), and observed strong upregulation of *Shh*, along with decreased *Frzb* expression, in *Pdzph1*-positive alpha2 tanycytes at P45 in TKO mice (Fig. 2F).

To infer cell lineage relationships between specific tanycyte subtypes and tanycyte-derived neural progenitors, we conducted RNA velocity analysis (*28*) on the full aggregated scRNA-Seq dataset (Fig. 2G). We observe that alpha2 tanycytes give rise to proliferating tanycytes, which in turn give rise to neural precursors following cell cycle exit (Fig. 2G, insets). It is noteworthy here that astrocytes appear to arise directly from alpha1 and alpha2 tanycytes, without going through a clear proliferative stage (Fig. 2G,H).

We then used pseudo-time analysis to identify six major temporally dynamic patterns of gene expression that occurred during the process of alpha2 tanycyte-derived neurogenesis (Fig. 2I, Table ST5). Based on the GO term enriched in each cluster (Fig. 2J), the transition from a quiescent to an actively proliferating state is associated with downregulation of metabolic genes (*Glul*), ion channels (*Kcnj10*), transcription factors (*Lhx2*) and Notch pathway components (*Notch1*) -- all of which are expressed at high levels in mature tanycytes (*3*, *10*). In addition, genes regulating ciliogenesis (*Dnah7a, Cfap65*) are rapidly downregulated. Following upregulation of genes controlling cell cycle progression and DNA replication (*Cenpf, Mcm3*), and cell cycle exit (*Btg2*), tanycyte-derived neural precursors upregulate genes that control chromatin conformation (*Phf3*), RNA splicing (*Sf3b1*), and neurogenesis (*Hes6*). This upregulation is then followed by expression of transcription factors that control specification of specific hypothalamic neuronal subtypes (*Dlx1, Lhx6*) and regulators of synaptogenesis (*Erbb4*), neurotransmitter biogenesis and reuptake (*Gad1, Pdyn, Slc32a1*), neurotransmitter receptors (*Grin1, Gria2*), and leptin signaling (*Lepr*).

To investigate changes of chromatin accessibility in TKO mice, we conducted scATAC-Seq analysis in FACS-isolated GFP-positive tanycytes and tanycyte-derived cells in both control and TKO mice at P8. UMAP analysis indicated that cell identity in both control and mutant samples could be readily assigned, based on gene expression data obtained from scRNA-Seq (Fig. 3A). The overall distribution of cell types was much like that seen for scRNA-Seq data (Fig. 2A), with more proliferating tanycytes and tanycyte-derived neurons observed in TKO cells compared to controls (Fig. 3B). Accessibility of the consensus NFI motif was assessed in all cell types in *Nfia/b/x*-deficient mice (Fig. 3C), and reduced levels of bound transcription factors were observed at these sites by footprinting analysis (Fig. 3E), indicating that Nfia/b/x are actively required to maintain accessible chromatin at a subset of their target sites.

**Figure 3:**
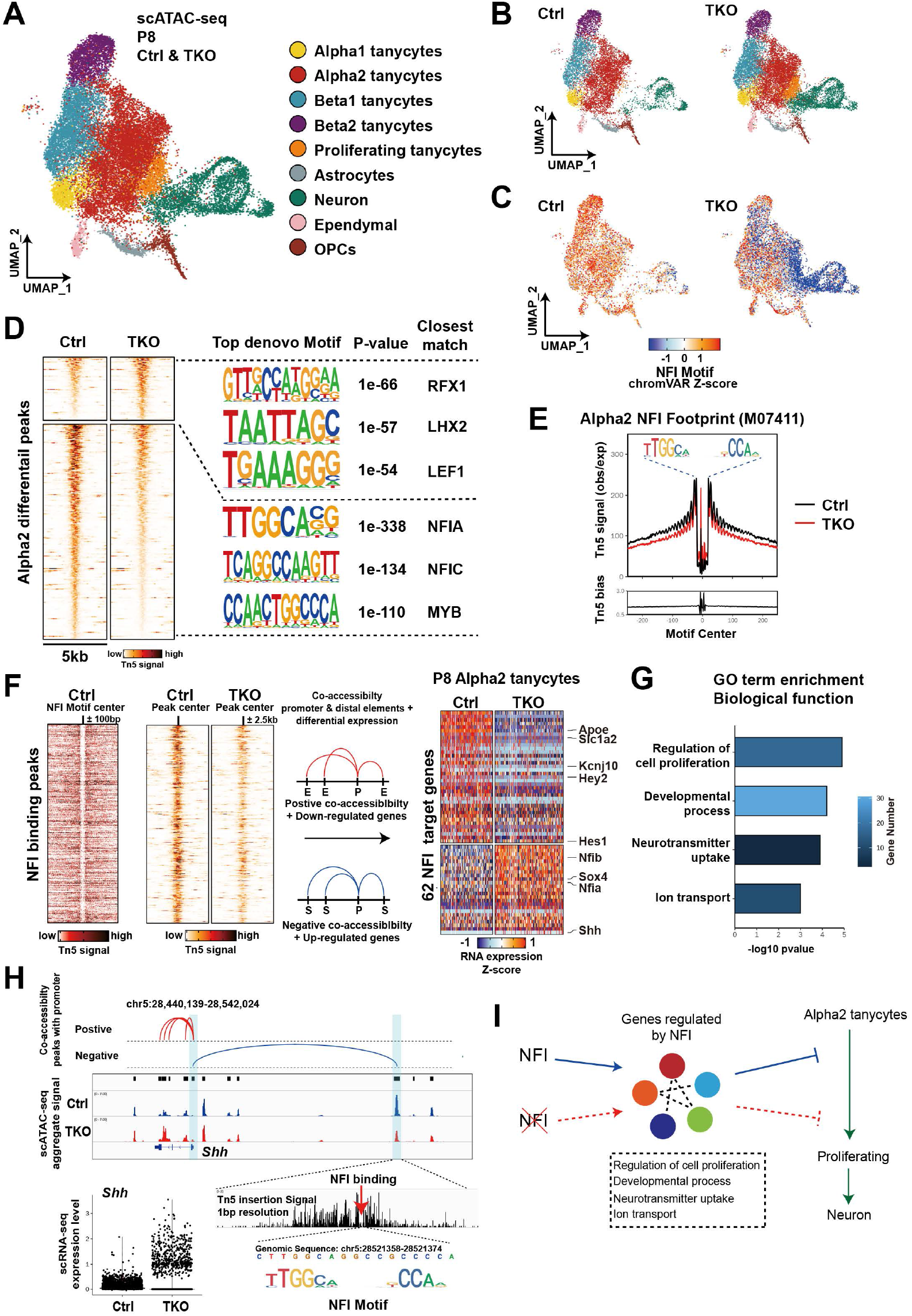
Single-cell ATAC-Seq analysis of P8 wildtype and *Nfia/b/x*-deficient tanycytes demonstrates derepression of Shh and increased Wnt signaling. A. Aggregate UMAP plot of scATAC-Seq data from control and NFI-deficient GFP+ tanycytes and tanycyte-derived cells isolated at P8. Cell types are indicated by color shading. B. Distribution of cell types shown for control and *Nfia/b/x*-deficient GFP+ cells at P8. C. Distribution of accessible consensus NFI motif shown for control and *Nfia/b/x*-deficient GFP+ cells. D. Transcription factor binding motifs selectively enriched and depleted in control and *Nfia/b/x*-deficient alpha2 tanycytes. E. Consensus NFI footprint distribution in control and *Nfia/b/x*-deficient alpha2 tanycytes. F. Integration of scATAC-Seq and scRNA-Seq data to identify differentially expressed genes in alpha2 tanycytes that are directly regulated by Nfia/b/x. G. Gene Ontology analysis of Nfia/b/x-regulated genes expressed in alpha2 tanycytes. H. *Shh* is directly repressed by Nfia/b/x in alpha2 tanycytes. I. Summary of Nfia/b/x action in alpha2 tanycytes.

We observed 639 chromatin regions that showed increased, and 3072 regions that showed decreased, accessibility in *Nfia/b/x*-deficient alpha 2 tanycytes relative to controls (Fig. 3D). As expected, HOMER analysis indicated that open chromatin regions (OCRs) specific to controls were most highly enriched for consensus sites for NFI family members (Fig. 3D, Table ST6). In contrast, motifs for the Wnt effector *Lef1* were enriched at OCRs specifically detected in *Nfia/b/x*-deficient alpha2 tanycytes. We observed that a subset of genes with altered expression in scRNA-Seq showed altered accessibility in putative associated cis-regulatory sequences, although changes in gene expression and chromatin accessibility often diverged (Table ST7, ST8). While putative regulatory elements of the downregulated genes *Aqp4, Hes1*, and *Fgf18* showed reduced accessibility, elements associated with other downregulated genes such as *Tgfb2* and *Sox8* showed increased accessibility. Likewise, while most upregulated genes showed increased accessibility, such as *Shh* and *Sox4*, some showed decreased accessibility. These divergent responses indicate that NFI genes control expression of a large number of transcription factors in tanycytes, which appear to perform dual roles as both activators and repressors.

To identify direct targets of NFI factors, and to better clarify their function in regulating proliferation and neurogenesis in alpha2 tanycytes, we integrated scRNA- and scATAC-Seq data from alpha2 tanycytes to identify genes with both altered expression and altered accessibility at sites containing NFI consensus sequences, and identified 62 genes in total (Fig. 3F, Table ST9). These include downregulated genes such as *Kcnj10, Apoe* and Notch pathway effectors such as *Hes1* and *Hey2*, as well as *Shh* and *Sox4*, which are upregulated, with direct target genes enriched for genes controlling proliferation and neural development (Fig. 3G). Transcriptions of Nfia and Nfib are themselves strongly activated by Nfia/b/x, consistent with findings in retina (*14*, *15*)). Importantly, we found NFI binding sites in peaks which are negatively correlated with the promoter of *Shh*, suggesting that NFI may directly repress *Shh* expression (Fig. 3H). Thus, NFI factors may act as both activators and repressors in alpha2 tanycytes that promote quiescence while inhibiting proliferation and neurogenesis (Fig. 3I).

### Shh and Wnt signaling regulate tanycyte proliferation and neurogenic competence

The increased expression of *Shh* and Wnt regulators that is observed in *Nfia/b/x*-deficient alpha2 tanycytes (Fig. 2E, Table ST4, Fig. 4A,B) suggested that Shh and Wnt signaling might promote proliferation and/or neurogenesis in tanycytes. We observe substantially increased expression of *Shh* in both alpha2 and beta1 tanycytes (Fig. 2F, S5), and more complex regulation of Wnt signaling modulators. We observe increased expression of *Sulf1*, which by regulating synthesis of heparan sulfate proteoglycans, typically enhances Wnt signaling (*29*). However, the broad-spectrum Wnt inhibitor *Notum*, which was recently shown to regulate quiescence in adult neural stem cells of the lateral ventricles (*30*), is upregulated from P17, potentially acting in a cell-autonomous manner counteracting the effects of increased cellular levels of Wnt signaling (Fig. 4A, B).

**Figure 4:**
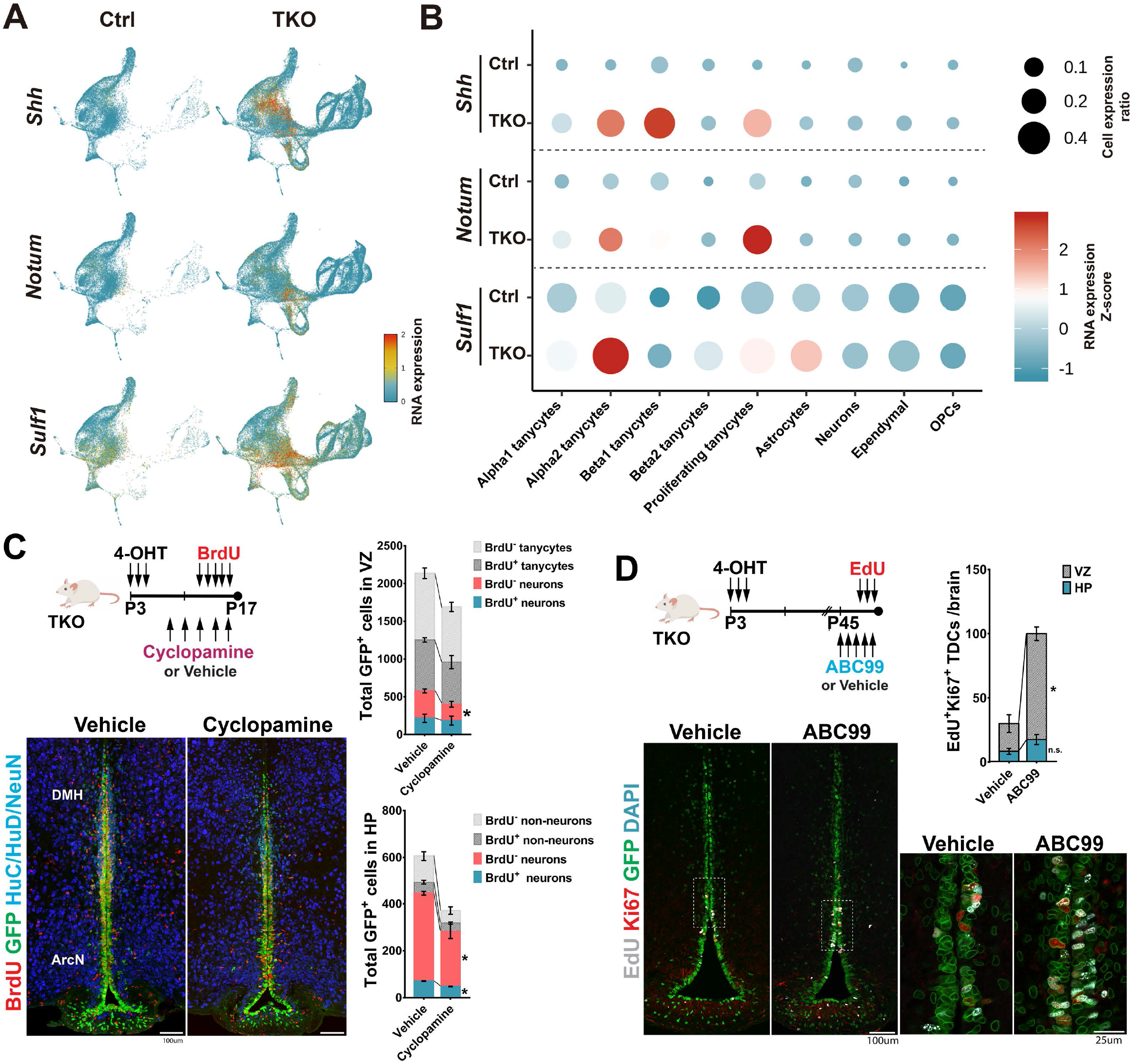
Shh and Wnt signaling stimulate proliferation and neurogenesis in *Nfia/b/x*-deficient tanycytes. A. Expression of *Shh*, *Notum* and *Sulf1* on aggregate UMAP plot. B. Dot plot showing expression level of Shh and Wnt pathway genes in each cluster. C. Shh inhibition by intraperitoneal (i.p.) cyclopamine inhibits neurogenesis in *Nfia/b/x*-deficient mice. D. Activation of Wnt signaling by inhibition of Notum via i.p. ABC99 induces proliferation of alpha tanycytes, and increased numbers of GFP+ proliferating cells in VZ and HP. VZ, ventricular zone; HP, hypothalamic parenchyma; TDCs, tanycyte-derived cells. Scale bars: C,D=100 μm; D(insets)=25 μm.

To determine whether this increased Shh inhibition might inhibit regulated tanycyte proliferation and/or tanycyte-derived neurogenesis, we administered the blood-brain barrier-permeable Shh antagonist cyclopamine via i.p. injection to *Nfia/b/x*-deficient mice every 2 days from P8 until P16, in conjunction with daily i.p. injections of BrdU from P12 to P16 (Fig. 4C). At these ages, Shh is both highly expressed in tanycytes and levels of proliferation and neurogenesis are high in *Nfia/b/x*-deficient alpha2 tanycytes. Cyclopamine administration resulted in a significant reduction in both the numbers of total GFP-positive cells and GFP/NeuN double-positive neurons in both the tanycytic layer in ventricular zone (VZ) and hypothalamic parenchyma (HP), when compared to vehicle controls, while BrdU incorporation was only significantly altered in parenchymal neurons, indicating a stronger effect on tanycyte-derived neurogenesis than on self-renewing tanycyte proliferation (Fig. 4C).

To determine whether Notum-dependent Wnt signaling played a role in inhibiting tanycyte proliferation and/or neurogenesis at later ages, we treated P45 *Nfia/b/x*-deficient mice with the blood-brain barrier-permeable Notum inhibitor ABC99 (*31*) once daily for 5 days, with EdU co-administered on the last 3 days. At this age, Notum expression is high and levels of tanycyte proliferation are substantially reduced relative to the early postnatal period. This led to a significant increase in proliferation in alpha2 tanycytes (Fig. 4D), indicating that activation of Wnt signaling stimulates tanycyte proliferation at later ages.

### *Nfia/b/x*-deficient tanycytes give rise to a diverse range of hypothalamic neuronal subtypes

To investigate the identity of tanycyte-derived neurons (TDNs), we analyzed a neuronal subset of scRNA-Seq data obtained from both control and *Nfia/b/x*-deficient mice (Fig. 5A). A total of 582 neurons were obtained from controls, while 15,489 neurons were obtained from *Nfia/b/x*-deficient mice (Table ST2). The great majority of control tanycyte-derived neurons were obtained at P8, while large numbers of tanycyte-derived neurons were seen at all ages in *Nfia/b/x*-deficient mice. UMAP analysis revealed that both control and *Nfia/b/x*-deficient tanycyte-derived neurons fell into two major clusters each of glutamatergic and GABAergic subtypes, with additional clusters corresponding to *Ascl1*/*Hes5*-positive postmitotic neural progenitor cells and immature *Dlx1/Sox11*-positive GABAergic precursors (Fig. 5A, B). RNA velocity analysis indicated three distinct major differentiation trajectories in both control and *Nfia/b/x*-deficient tanycyte-derived neurons, which give rise to the two major glutamatergic clusters and the GABAergic neurons (Fig. 5C).

**Figure 5:**
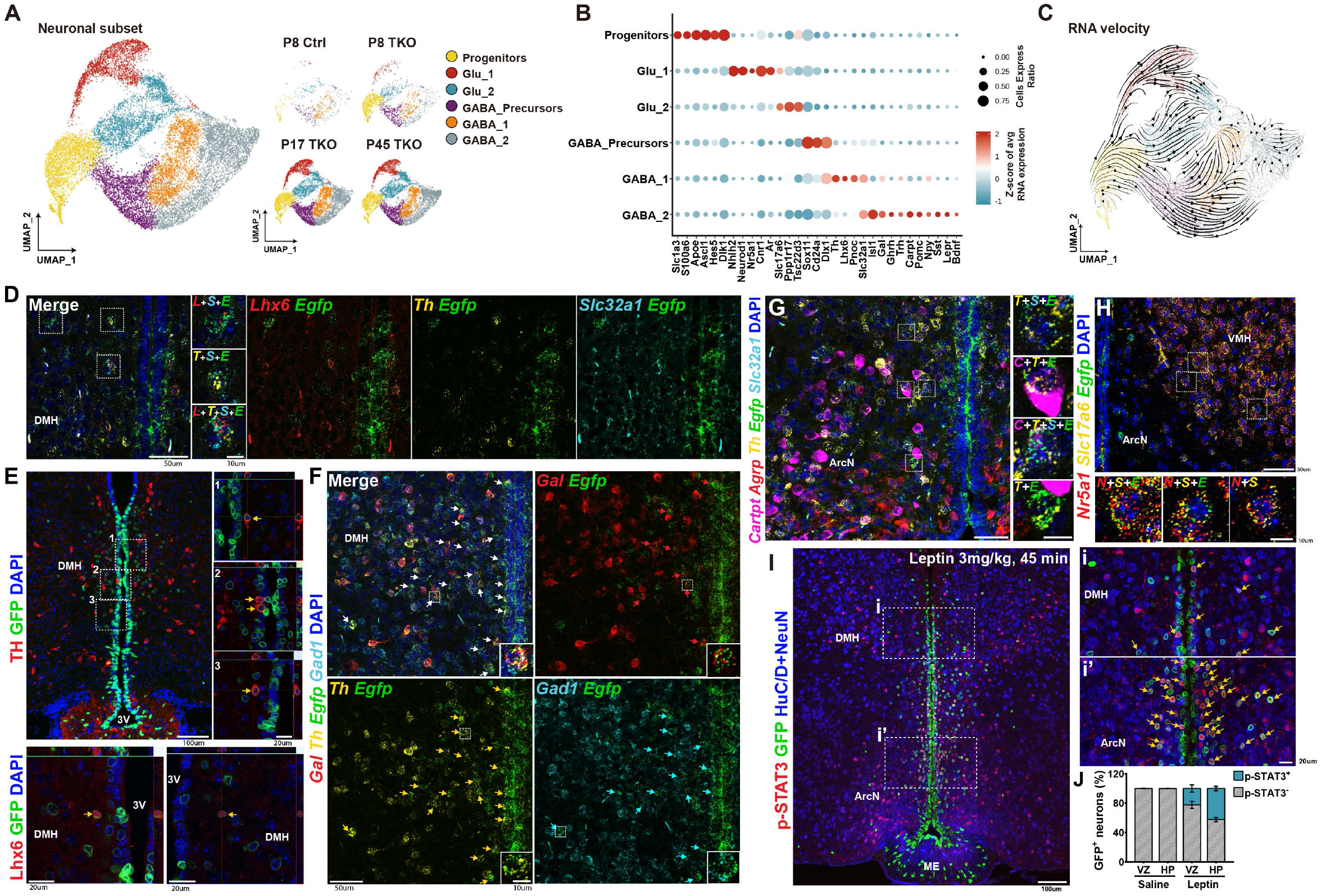
Identification of selective markers of tanycyte-derived neurons. A. UMAP plot showing major clusters of tanycyte-derived neuronal subset, separated by age and genotype. B. Dot plot showing major subtype-specific markers of tanycyte-derived neurons. C. RNA velocity analysis indicates differentiation trajectories for tanycyte-derived neurons. D. SmfISH analysis and E. immunohistochemistry demonstrates expression of *Th* and *Lhx6* in *Egfp+* tanycyte-derived neurons in *Nfia/b/x*-deficient mice. F. SmfISH analysis of *Gal, Gad1* and *Th* in *Egfp+* tanycyte-derived neurons in *Nfia/b/x*-deficient mice. G. *Cart, Agrp, Slc32a1* and *Th* expression in *Egfp+* tanycyte-derived neurons in *Nfia/b/x*-deficient mice. H. *Nr5a1* and *Slc17a6* expression in *Egfp+* tanycyte-derived neurons in *Nfia/b/x*-deficient mice. All insets are enlarged images of examples of colocalization (white boxes in D,E,F,G,H). I. pStat3 staining 45 minutes after i.p. administration of 3 mg/kg leptin in *Nfia/b/x*-deficient mice (n=3 mice). Arrows indicate GFP+/pStat3+ tanycyte-derived neurons. Insets show higher magnification images in DMH (i) and ArcN (i’). J. Fraction of pStat3-positive tanycyte-derived neurons in VZ and HP after leptin administration. pStat3 was not induced in saline-injected mice (n=2 mice). VZ, ventricular zone; HP, hypothalamic parenchyma. Scale bars: D, F, G, H=50 μm; insets in D, F, G, H=10 μm; E, H, I=100 μm; high magnification images in E =20 μm.

53% of tanycyte-derived neurons were GABAergic -- as determined by expression of *Gad1, Gad2* and/or *Slc32a1* -- while 30% were *Slc17a6*-positive glutamatergic neurons (Fig. 5B, Table ST10, ST11). Glu_1 was enriched for the transcription factors *Nhlh2* and *Neurod2*, as well as markers of glutamatergic VMH neurons, such as *Nr5a1, Cnr1*, and the androgen receptor *Ar*, while Glu_2 was enriched for markers of glutamatergic DMH neurons, such as *Ppp1r17*, and ArcN markers such as *Chgb* (Fig. 5B). GABAergic neurons expressed a diverse collection of molecular markers expressed by neurons in the ArcN and DMH, as well as the adjacent zona incerta (ZI), which regulates a broad range of internal behaviors including feeding, sleep and defensive behaviors (*32*). GABA_1 was enriched for a subset of ZI and DMH-enriched genes (*Lhx6, Pnoc*), while GABA_2 was enriched for genes selectively expressed in ArcN neurons (*Isl1*), as well as genes expressed in GABAergic neurons in both the ArcN and DMH (*Cartpt, Npy, Sst, Gal, Trh, Th*).

We used both multiplexed smfISH (Fig. 5D,F) and immunohistochemistry (Fig. 5E, ST12) to confirm expression of *Th*, *Lhx6*, and *Gal* in GFP-positive *Nfia/b/x*-deficient tanycyte-derived GABAergic neurons in the dorsomedial hypothalamus, as verified by *Egfp/Slc32a1 (or Gad1*) co-labeling. Expression of *Cartpt* and *Th* was observed in GABAergic tanycyte-derived neurons in both the ArcN and DMH (Fig. 5G, S6A), and coexpression of Lhx6 and Th, as well as Cartpt and Th, was observed using immunohistochemistry in a subset of tanycyte-derived and non-tanycyte-derived neurons. Small numbers of *Nr5a1*-expressing glutamatergic (*Slc17a6*-positive) neurons were detected in the dorsomedial VMH (Fig. 5H). No expression of markers specific to neurons of more anterior or posterior hypothalamic regions (e.g. *Avp, Crh, Oxt, Pmch, Hcrt, Vip*, etc) was detected (Table ST11).

These data demonstrate that tanycyte-derived neurons express molecular markers of multiple neuronal subtypes located in the tuberal hypothalamus, including the ArcN and VMH. To determine how closely the gene expression profile of tanycyte-derived neurons more broadly resembles the profile of hypothalamic neurons in these regions, we used LIGER analysis (*33*) to integrate clustered scRNA-Seq from a previous study of adult ArcN, in which a small number of VMH neurons was also profiled (*23*) (Fig. S6B, Table ST13). Integration of these datasets using LIGER (*34*), and comparison of cell types in each cluster using alluvial plotting (Fig. S6C), indicate substantial overlap in cellular gene expression profiles between glutamatergic tanycyte-derived neurons cluster Glu_1 with both *Kiss1/Tac2-positive* ArcN neurons and *Nr5a1*-positive VMH neurons. Glu_2 overlapped with *Rgs16/ Nmu*-positive ArcN neurons. GABAergic tanycyte-derived GABA_1 and GABA_2 clusters overlapped with several different ArcN neuronal clusters, including clusters that contained neurons expressing *Th*, *Ghrh* and/or *Trh*. In contrast, some subtypes of ArcN neurons were represented only sparsely or not at all among tanycyte-derived neurons, while other tanycyte-derived neurons appeared to correspond to cell types not found in the published scRNA-Seq dataset. For instance, while some tanycyte-derived neurons closely resembled *Pomc*-expressing ArcN neurons, the relative fraction of *Pomc*-positive tanycyte-derived neurons was substantially lower than in ArcN (LIGER Cluster 4). Likewise, no tanycyte-derived neurons mapped to LIGER Cluster 7, which corresponded to *Agrp*-positive ArcN neurons, and no *Agrp*-positive tanycyte-derived neurons were detected using smfISH (Fig 5G, S6B-C). While few immature tanycyte-derived neurons mapped to neurons in the mature ArcN scRNA-Seq dataset, as expected, two clusters of mature glutamatergic (LIGER Clusters 8 and 9) and one cluster of GABAergic (LIGER Cluster 11) also showed little correspondence to ArcN neurons, and appeared to correspond to DMH-like neurons based on their expression of DMH-enriched genes (e.g. *Ppp1r17, Lhx6*).

At P45, 4.4% of tanycyte-derived neurons in the GABA_2 cluster expressed the leptin receptor *Lepr* (Fig. 5B), despite *Lepr* being essentially undetectable in tanycytes themselves, as previously reported (*19*). As a result, we tested whether tanycyte-derived neurons were capable of responding to leptin signaling. P90 mice that had undergone an overnight fast were injected i.p. with 3mg/kg leptin, and sacrificed after 45 minutes, providing sufficient time for leptin-responsive neurons throughout the hypothalamus to induce pStat3 immunoreactivity (*35*, *36*). We observed robust induction of pStat3 immunoreactivity in GFP-positive tanycyte-derived neurons under these conditions (Fig. 5I), with low levels of immunoreactivity under unstimulated conditions, as previously reported (*35*, *36*). A total of 42.3 (± 2.5) % of parenchymal tanycyte-derived neurons in DMH and ArcN show pStat3 induction under these conditions, along with 22.3 (± 3.9) % of tanycyte-derived neurons in the subventricular region (Fig. 5J), confirming that a subset of tanycyte-derived neurons are leptin-responsive.

### Neurons derived from *Nfia/b/x*-deficient tanycytes integrate into hypothalamic neural circuitry

Since our scRNA-Seq analysis identified the majority of tanycyte-derived cells as neurons in *Nfia/b/x*-deficient mice, we next investigated whether these cells showed electrophysiological properties of functional neurons. To characterize tanycyte-derived cells, we performed whole-cell patch-clamp recordings from GFP-positive parenchymal cells in acute brain slices obtained from *Nfia/b/*x-deficient mice that had undergone 4-OHT treatment between P3 and P5. Biocytin filling of recorded tanycyte-derived cells revealed neuron-like morphology, typically showing 3-5 major dendritic processes, similar to GFP-negative control neurons (Fig. 6A, Fig. S7). We found that the majority of GFP-positive tanycyte-derived cells in the hypothalamic parenchyma fired action potentials in response to depolarizing current steps (Fig. 6B,C, 95%, 40 of 42 cells from P15-P97). In contrast, GFP-positive cells located in the tanycytic layer retained nonspiking, glial-like electrophysiological properties in *Nfia/b/x*-deficient mice (Fig. S8).

**Figure 6:**
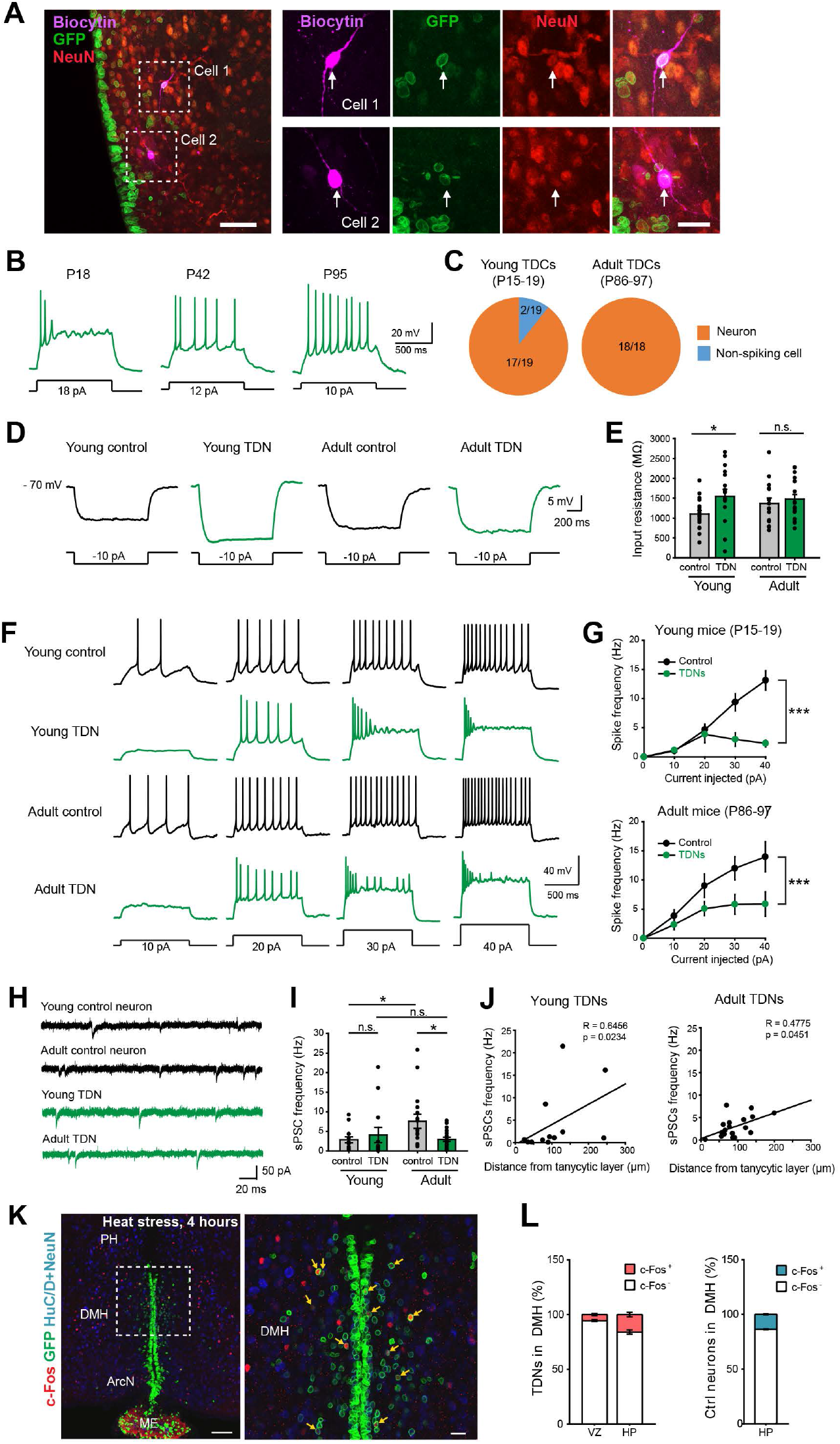
*Nfia/b/x*-deficient tanycytes differentiate into neurons, integrate into hypothalamic neural circuitry, and respond to physiological stimuli. A. Low and high magnification confocal images showing two biocytin-filled GFP+ recorded cells (white arrows) in an NFI TKO brain slice stained with NeuN. B. Example responses of tanycyte-derived cells to depolarizing current steps. C. The proportion of tanycyte-derived neurons among tested tanycyte-derived cells in young (left) and adult (right) mice. D. Representative average responses to hyperpolarizing current steps. E. Summary graphs of input resistance (young control neurons, 18 cells from 5 mice, 1,101 ± 89 MΩ, young tanycyte-derived neurons, 17 cells from 5 mice, 1,546 ± 176 MΩ, p = 0.0238, Mann-Whitney U test; adult control neurons, 16 cells from 6 mice, 1,369 ± 132 MΩ, adult tanycyte-derived neurons, 18 cells from 6 mice, 1,477 ± 119 MΩ, p = 0.4777, Mann-Whitney U test). F. Representative voltage traces recorded from control and tanycyte-derived neurons from young and adult mice evoked by 10-40 pA depolarizing current steps as indicated. G. The current-spike frequency relationships measured from control and tanycyte-derived neurons from young mice (top) and adult mice (bottom). The current-frequency was significantly different between tanycyte-derived and control neurons at both ages (Young mice: control neurons, 14 cells from 4 mice; tanycyte-derived neurons, 13 cells from 4 mice, p <0.0001, two-way ANOVA; Adult mice: control neurons, 16 cells from 6 mice, tanycyte-derived neurons, 18 cells from 6 mice, p = 0.0001, two-way ANOVA). H. Representative traces of spontaneous postsynaptic currents (sPSCs). I. Summary graphs of sPSC frequency (young control neurons, 14 cells from 4 mice, 2.85 ± 0.74 Hz, young tanycyte-derived neurons, 13 cells from 4 mice, 4.07 ± 1.93 Hz, p = 0.2983, Mann-Whitney U test; adult control neurons, 16 cells from 6 mice, 7.56 ± 1.82 Hz, adult tanycyte-derived neurons, 18 cells from 6 mice, 2.98 ± 0.54 Hz, p = 0.0324, Mann-Whitney U test). J. Positive correlation between the distance from the tanycytic layer for each tanycyte-derived neuron and its sPSC frequency in both young (left, 12 cells from 4 mice, p = 0.0234, Spearman’s Rho correlation) and adult (right, 18 cells from 6 mice, p = 0.0451, Spearman’s Rho correlation) mice. K. 4 hr heat stress (38°C) selectively induced c-fos expression in tanycyte-derived neurons in DMH (higher magnification inset shown in right). L. Fraction of c-fos-positive tanycyte-derived neurons in VZ and HP of DMH, and fraction of all c-fos-positive and negative neurons in DMH (n=3 mice). ArcN=arcuate nucleus, DMH=dorsomedial hypothalamus, HP=hypothalamic parenchyma, ME=median eminence, PH=posterior hypothalamus, VZ=ventricular zone. Scale bars: A. Left=50 μm, right 20 μm, K=100 μm (inset=20 μm).

Like typical hypothalamic neurons (*37*), many tanycyte-derived neurons fired spontaneous action potentials (sAPs) and exhibited relatively depolarized resting membrane potentials (Fig. S9A). However, the proportion of tanycyte-derived neurons showing sAPs was significantly lower than that of GFP-negative control neurons in young (P15-17) mice (Fig. S9B). In addition, we found that the input resistance was significantly higher in tanycyte-derived neurons than in GFP-negative control neurons in young (P15-19) mice, although the input resistances were similar for tanycyte-derived and control neurons in adult mice (P86-P97) (Fig. 6D,E, Table ST14). Together, these data indicate that, although there were some differences with GFP-negative control neurons, almost all tanycyte-derived cells fired action potentials and shared electrophysiological features of control neurons, indicating that they are indeed neurons.

Since the great majority of tanycyte-derived cells appear to be functional neurons, we next asked whether their evoked action potential firing properties were similar to those of neighboring GFP-negative hypothalamic neurons. Although tanycyte-derived neurons fired action potentials to depolarizing current steps, the average current-frequency curve of tanycyte-derived neurons was significantly different from the curve for control neurons in both young and adult mice (Fig. 6F,G). Although the number of action potentials elicited initially increased with age, the tanycyte-derived neurons were unable to reliably generate repetitive actions potentials with larger current steps in contrast to control neurons, leading to saturation of the current-frequency curves (Fig. 6G; Fig. S9C,D). These results suggest that tanycyte-derived neurons may have a different ion channel composition than control neurons and may not be as developmentally mature.

We next asked whether tanycyte-derived neurons receive synaptic inputs from other neurons and are functionally integrated into hypothalamic circuits. We detected spontaneous postsynaptic currents (sPSCs) in 34 of 35 recorded tanycyte-derived neurons (Fig. 6H,I). However, the frequency of sPSCs in adult tanycyte-derived neurons was significantly lower than for control neurons in adult mice (Fig. 6H, I Table ST14), suggesting that the number of functional synapses received is nonetheless fewer than for wild type neurons. As our histological data suggest that tanycyte-derived neurons undergo progressive radial migration away from the tanycyte layer into the hypothalamic parenchyma as they mature, we asked whether there was any correlation between a tanycyte-derived neuron’s distance from the tanycyte layer and the number of synaptic inputs it receives. We observed a positive correlation between a tanycyte-derived neuron’s distance from the tanycyte layer and the sPSC frequency in both young and adult mice, suggesting that the farther a tanycyte-derived neuron migrates from the tanycytic layer, the more functional synapses from local neurons it receives (Fig. 6J).

Having established that tanycyte-derived neurons integrate into hypothalamic circuits, we next tested whether tanycyte-derived neurons are activated in response to changes in internal states that modulate the activity of nearby hypothalamic neurons. We first investigated the response of tanycyte-derived neurons in the DMH to heat stress, which is known to robustly modulate the activity of DMH neurons (*38*, *39*). At P45, we observed that 16.0 ± 1.6% of parenchymal tanycyte-derived neurons in the DMH induce c-fos expression following 4 hrs of exposure to 38°C ambient temperature in *Nfia/b/x*-deficient mice, which is essentially equivalent to the portion of GFP-negative control neurons activated, 13.7± 0.3% (Fig. 6K,L).

## Discussion

This study demonstrates that tanycytes retain the ability to generate a broad range of different subtypes of hypothalamic neurons in the postnatal brain, and that this latent ability is actively repressed by NFI family transcription factors. Induction of proliferative and neurogenic competence by selective loss of function of *Nfia/b/x* leads to the robust generation of hypothalamic neuronal precursors that undergo outward radial migration, mature, and integrate into existing hypothalamic circuitry. Tanycyte-derived neurons respond to dietary signals, such as leptin, and heat stress. This implies that tanycyte-derived neurogenesis can potentially modulate a broad range of hypothalamic-regulated physiological processes.

NFI factors have historically been primarily studied in the context of promoting astrocyte specification and differentiation (*40*, *41*), and loss of function of *Nfia/b/x* disrupts generation of tanycyte-derived astrocytes, ependymal cells and oligodendrocyte progenitors, and downregulates expression of tanycyte-enriched genes that are also expressed in astrocytes such as *Kcnj10* and *Aqp4* (Fig. 2). In addition to their role in promoting gliogenesis, however, recent studies have shown that NFI factors confer late-stage temporal identity on retinal progenitors (*14*, *42*), allowing generation of late-born bipolar neurons and Müller glia, and decreasing proliferative and neurogenic competence. Selective loss of function of *Nfia/b/x* in mature Müller glia likewise induces proliferation and generation of inner retinal neurons (*15*), although the levels seen are lower than seen following loss of function in retinal progenitors (*14*). As in retina, *Nfia/b/x* are more strongly expressed in late than early-stage mediobasal hypothalamic progenitors (*16*), and adult tanycytes show substantially lower levels of proliferation following loss of function of *Nfia/b/x* than is seen in neonates (Fig. 1). However, the levels of proliferation and neurogenesis seen in neonatal tanycytes are much greater than those seen in Müller glia, which likely reflects the fact that mammalian Müller glia proliferate only rarely and essentially lack neurogenic competence (*11*), while tanycytes retain limited neurogenic competence (*9*). This implies that NFI factors may be part of a common gene regulatory network that represses proliferation and neurogenic competence in radial glia of the postnatal forebrain and retina.

Using scRNA and ATAC-Seq analysis, we identified multiple genes that are strong candidate extrinsic and intrinsic regulators of proliferative and neurogenic competence in tanycytes. We observe that loss of function of *Nfia/b/x* effectively regresses tanycytes to a progenitor-like state. Both Shh (*43*–*45*) and Wnt (*46*–*48*) signaling are required for progenitor proliferation and neurogenesis in embryonic tuberal hypothalamus, and our study demonstrates that they play similar roles in *Nfia/b/x*-deficient tanycytes. Tanycyte-specific loss of function of *Nfia/b/x* both downregulates Notch pathway components and upregulates the Notch inhibitor *Dlk1*, while also downregulating *Tgfb2*. Both Notch signaling and *Tgfb2* promote quiescence and inhibit proliferation in retinal Müller glia and cortical astrocytes (*49*–*52*), and likely play a similar role in tanycytes.

*Nfia/b/x*-deficient tanycytes likewise downregulate transcription factors that are required for specification of astrocytes and Müller glia – including *Sox8/9 (53, 54*) – while upregulating neurogenic bHLH factors such as *Ascl1* and *Sox4. Ascl1* is both required for differentiation of VMH neurons (*55*) and also sufficient to confer neurogenic competence on retinal Müller glia (*56*, *57*). NFI factors thus control expression of a complex network of extrinsic and intrinsic factors that regulate neurogenic competence in tanycytes, and it may be possible to further stimulate tanycyte-derived neurogenesis by modulating select components of this network.

scRNA-Seq analysis reveals that tanycyte-derived neurons arise from *Ascl1+* precursors and are heterogeneous, falling into several molecularly distinct clusters. The gene expression profiles of control and *Nfia/b/x*-deficient tanycyte-derived neurons closely resemble one another, indicating that NFI factors are not obviously required for differentiation of individual neuronal subtypes, in contrast to their role in retina and cerebellum (*14*, *58*, *59*). However, the number of tanycyte-derived neurons in control samples drops dramatically after P8, with few detected at P45 (Fig. 2). We observe an age-dependent decline in the number of tanycyte-derived neurons in controls, potentially in line with the high levels of cell death reported in postnatally generated hippocampal and olfactory bulb neurons (*60*). However, this decline is insufficient to account for this effect, and it may instead result from the well-established difficulties in obtaining viable, dissociated mature neurons following FACS for whole-cell scRNA-Seq analysis (*61*).

Tanycyte-derived neurons are predominantly found in the DMH and ArcN, with much smaller numbers detected in the VMH and ME (Fig. 1H). They are mostly GABAergic, and express molecular markers of DMH and ArcN neurons (Fig. 5A,B), and substantial subsets closely match scRNA-Seq profiles of neuronal subtypes obtained from ArcN and VMH (Fig. S6) (*23*). They include neuronal subtypes that regulate feeding, sleep, and directly regulate pituitary function, and many other subtypes whose function has yet to be characterized. In light of findings that tanycyte-derived neurogenesis can be stimulated by dietary and hormonal signals, and potentially modulate body weight and activity levels (*3*, *7*), this raises the possibility that different internal states may trigger generation and/or survival of functionally distinct tanycyte-derived neuronal subtypes, leading to long-term changes in both hypothalamic neural circuitry and physiological function.

We observe that tanycyte-derived neurons survive for months and integrate into hypothalamic neuronal circuitry, with subsets showing c-fos induction in response to heat stress (Fig 6). Older tanycyte-derived neurons receive more synaptic inputs than younger tanycyte-derived neurons, and their input resistance lowers to become equivalent to that of GFP-negative nearby neurons, demonstrating progressive maturation. However, the frequencies of spontaneous postsynaptic currents and the evoked action potentials of tanycyte-derived neurons remains consistently lower than those of GFP-negative control neurons. This may be an intrinsic property of tanycyte-derived neurons, distinguishing them with pre-existing local neurons. Alternatively, the excess tanycyte-derived neurons generated from *Nfia/b/x*-deficient tanycytes may form synaptic connections less efficiently. Distinguishing these possibilities will require electrophysiological recording of tanycyte-derived neurons from control animals, although the far smaller population of these cells in wildtype animals make this experiment very challenging.

While levels of tanycyte-derived neurogenesis are low under baseline conditions, adding small numbers of newly generated neurons to existing hypothalamic neural circuits may have important effects on hypothalamic-regulated behaviors and physiology. Furthermore, selective loss of specific hypothalamic neuronal subtypes, and accompanying metabolic and behavior changes, is observed both in neurodegenerative disorders such as Alzheimer’s disease and frontotemporal dementia (*62*), and in conditions such as obesity and type 2 diabetes (*63*). Identifying new means of enhancing neurogenic competence in mature tanycytes, while selectively promoting differentiation of specific tanycyte-derived neuronal subtypes, may therefore ultimately prove useful in generating long-term changes in hypothalamic physiology in a variety of therapeutic contexts.

## Acknowledgements

We thank R. Gronostajski for providing *Nfia/b/x* conditional mutant mice. J. Ling, A. Kolodkin, J. Nathans, D. Lee, J, Bedont, M. Placzek and W. Yap for helpful comments on the manuscript. We thank the Hopkins Multiphoton Imaging Core for assistance with microscopy, and the St Jude Cloud facility for hosting the scRNA-Seq and scATAC-Seq data. This work was supported by grant DK108230 to S. Blackshaw, a Maryland Stem Cell Postdoctoral Research Fellowship to S. Yoo, and Imaging Core Grant P30NS050274.

## Data availability

All scRNA-Seq and scATAC-Seq data presented here are accessible at GEO (GSE160378), and can be queried interactively at https://viz.stjude.cloud/stjude/visualization/tanycyte.

## Methods and Materials

### Contact for Reagent and Resource Sharing

Further information and requests for resources and reagents should be directed to and will be fulfilled by the Lead Contact, Seth Blackshaw

### Animals

Rax-CreER^T2^ mice (The Jackson Laboratory Stock No. 025521) generated in the laboratory (*18*) were crossed with the Cre-inducible Sun1-GFP reporter (*20*) (*Sun1-sfGFP-Myc*, The Jackson Laboratory Stock No. 021039, provided by Jeremy Nathans). *Nfia^lox/lox^ (14, 15), Nfib^lox/lox^ (64*), and *Nfix^lox/lox^ (65*) mice were used to generate *Nfia/b/x* homozygous triple mutant mice previously described (*14*, *15*). To generate tanycyte-specific loss of function mutants of *Nfia/b/x* genes, the triple mutant mice were crossed to Rax-CreER^T2^; Sun1-GFP mice. To induce Cre recombination, these mice were intraperitoneally (i.p.) injected with 0.2 mg 4-hydroxytamoxifen (4-OHT) dissolved in corn oil for 3 consecutive days from postnatal day (P)3 to P5. Mice were housed on a 14 hr - 10 hr light/dark cycle (07:00 lights on – 21:00 lights off) in a climate-controlled pathogen-free facility. All experimental procedures were preapproved by the Institutional Animal Care and Use Committee (IACUC) of the Johns Hopkins University School of Medicine.

### BrdU and EdU incorporation assay

To label the proliferating cells, BrdU (Sigma #B5002) was dissolved in saline solution and 100 mg/kg of body weight was i.p. injected for 5 consecutive days for the dates indicated. For the AAV1-Cre-mCherry stereotaxic injection, an osmotic mini-pump (Alzet model 1002, #0004317) was filled with BrdU dissolved in aCSF (TOCRIS #3525) and installed immediately into the hole remaining after the virus injection needle was removed. The 2 cm long tube connecting the mini-pump and cannula was filled with aCSF, so that the actual 30 μg/day infusion of BrdU was started from the third day following implantation. To avoid potential toxic effects of long-term BrdU exposure, we used EdU (ThermoFisher #A10044) for quantitative studies of cell proliferation. For this purpose, we used a 50 μg/g dose of EdU, which has been previously validated for proliferation studies in the adult brain (*66*).

### Inhibitory drug administration

ABC99 (Sigma #SML2410) was prepared as previously described (*67*), except for the fact that a 16.5 mg/ml stock solution was used. This stock was sequentially mixed with Tween-80 (Sigma #P1754), PEG-400 (Merch #91893), and 0.9% NaCl in the ratio of 1:1:1:17. P45 *Nfia/b/x* triple knockout male mice were i.p. injected with 10 mg/kg ABC99 for 5 consecutive days. 50 mg/kg body weight EdU was injected together from the third day of treatment in order to profile proliferation induced by Notum inhibition (*30*). For the 1 μg/ul Cyclopamine stock solution, 1 mg Cyclopamine (TOCRIS, #1623) powder was dissolved into 1 ml 2-hydroxylpropyl-beta-cyclodextrin (Sigma #C0926) prepared as a 45% solution in phosphate buffered saline (PBS). 10 μg/g Cyclopamine was injected from P8 to P16 on alternative days. 100 μg/g body weight BrdU was injected together from the 3rd injection for 5 consecutive days. Control mice were treated with the corresponding vehicle solution.

### Tissue processing and immunohistochemistry

Mice were anesthetized with an i.p. injection of Tribromoethanol/Avertin, followed by transcardial perfusion with 2% paraformaldehyde (PFA) as previously described (*68*). Brains were dissected, postfixed in the same fixative, and prepared for the cryopreservation in O.C.T. embedding compound. A series of 25 μm coronal sections were stored in antifreeze solution at −20°C until ready for immunostaining.

After brief washing with 1X PBS to remove the antifreeze solution, sections were mounted on Superfrost Plus slides (Fisher Scientific) before starting immunohistochemistry, and dried at room temperature for 30 min. To ensure adequate fixation for nuclear staining, sections were immersed into the fixative solution for 10 min at this point. Antigen retrieval was performed by incubating the slides with the prewarmed sodium citrate buffer (10 mM sodium citrate, pH 6.0) in an 80°C water bath for 30 min. For HuC/D antibody staining, sections were also treated with 0.3% hydrogen peroxide in order to block endogenous peroxidase activity prior to the blocking step with 10% horse serum/0.1% Triton X-100 in 1X PBS for an hour. pSTAT3 staining required different pretreatment as described before (*19*, *68*). After finishing the first round of immunostaining, the fluorescence signal was crosslinked by incubation in 2% PFA for 10 min, followed by either EdU staining using the Click-iT EdU detection kit conjugated with Alexa Fluor 647 (ThermoFisher #C10340) or BrdU antibody staining. For BrdU staining, freshly prepared 2N HCl was spread on the slides and incubated at 37°C for 30 min on the humidified chamber. 0.1M Borate buffer (pH 8.5) was used for acid neutralization by incubating for 10 min at room temperature. Antibodies used were listed in Table ST1. After counter staining with DAPI, the slides were coverslipped with Vectashield antifade mounting medium (Vector Laboratories, #H-1200) and dried at room temperature for no more than 30 min. The slides were stored at 4°C and imaged within two days to achieve the best quality using a Zeiss LSM 700 Confocal at the Microscope Facility at Johns Hopkins University School of Medicine.

### RNAscope Hiplex assay

For the RNAscope Hiplex assay, P45 triple knock-out and control male mice were sacrificed by cervical dislocation and brains were dissected out. The brains were immediately immersed in 4% PFA in DEPC-treated 1X PBS and incubated overnight at 4°C. All other sample preparation procedures were performed as recommended in the manufacturer’s instructions for OCT-embedded fresh frozen tissue preparation. 14 μm sections were cut on a cryostat and briefly washed with 1X PBS before mounting on Superfrost Plus slides (Fisher Scientific). The slides were dried at −20°C and stored at −80°C before use. The Hiplex assay was performed by following the manufacturer’s instructions using probes listed in Table S12. The sections were imaged on a Zeiss LSM 800 Confocal at the Multiphoton Imaging Core in the Department of Neuroscience at Johns Hopkins University School of Medicine.

### Heat stimulation and leptin injection

P45 male mice were exposed to ambient heat (38°C) for 4 hours (*69*) by incubating in a prewarmed light-controlled cabinet in the rodent metabolism core facility at the Center for Metabolism and Obesity Research of the Johns Hopkins University School of Medicine. During the procedure, mice were provided ad libitum access to water and food and carefully monitored. Transcardiac perfusion with 4% PFA in 1X PBS was performed immediately after heat exposure. The dissected brains were processed as described above and used for c-fos immunostaining. Leptin injection was performed on P90 male mice that were fasted for 18 hours prior to treatment. 3 mg/kg body weight of leptin (PeproTech, #450-31) dissolved in saline solution was i.p. injected, and 45 min later transcardial perfusion was performed using 2% PFA as described above.

### Cell counting and statistical analysis

All cell counts were performed blindly and manually by five independent observers using Fiji/ImageJ software. Five sections corresponding to −1.55, −1.67, −1.79, −1.91, −2.15 mm from Bregma were chosen among the serial sections for cell counting. Initially, cell numbers were normalized by the size (mm) of hypothalamic nuclei measured. Because *Nfia/b/x*-deficient animals did not show any obvious structural differences, in subsequent experiments, we used absolute numbers. All values are expressed as mean ± S.E.M. Comparisons were analyzed by two-tailed Student’s t-test using Microsoft Excel unless stated otherwise. A p-value of < 0.05 was considered statistically significant.

### Cell preparation for scRNA-seq

P8, P17, and P45 *Rax-CreER;Nfia^lox/lox^;Nfib^lox/lox^;Nfix^lox/lox^;CAG-lsl-Sun1-GFP* mutant mice and control *Rax-CreER;CAG-lsl-Sun1-GFP* mice were sacrificed by cervical dislocation and brains were dissected. One biological replicate of each timepoint and genotype were analyzed, with the exception of P45 TKO, where two biological replicates were analyzed. 2 mm thick coronal slices including the hypothalamic protruding median eminence (ME) were collected using adult mouse brain matrix (Kent Scientific). The mediobasal hypothalamic region was microdissected using a surgical scalpel, dampened in Hibernate-A media supplemented with 0.5 mM GlutaMax and 2% B-27 (HABG), and chopped with a razor blade. Brain tissues were transferred into preequilibrated papain/Dnase-I mix (Worthington Papain Dissociation System, #LK003150) and incubated for 30 min at 37°C with frequent agitation using a fire-polished glass pipette. Dissociated cells were filtered through a 40 μm strainer and subjected to density gradient centrifugation to remove the cell debris as suggested in the manufacturer’s protocol. Cells were resuspended in HABG medium and GFP-positive cells were FACS isolated in the Bloomberg Flow Cytometry and Immunology Core at Johns Hopkins University. Cells were resuspended with ice-cold PBS containing 0.04% BSA and 0.5 U/μl RNase inhibitor, and 10,000-15,000 cells were loaded on a 10x Genomics Chromium Single Cell system (10X Genomics, Redwood City, CA), using the v3 chemistry following the manufacturer’s instructions, and libraries were sequenced on an Illumina NextSeq with ~200 million reads per library. Sequencing results were processed through the Cell Ranger 3.1 pipeline (10x Genomics, Redwood City, CA) with default parameters.

### Single-cell ATAC-Seq

Single cell ATAC-Seq was performed using the 10x Genomic single cell ATAC reagent V1 kit following the manufacturer’s instructions. Briefly, FACS-sorted cells (~30k cells) were centrifuged at 300xg for 5 min at 4°C. The cell pellet was resuspended in 100 μl of Lysis buffer, mixed 10x by pipetting and incubated on ice for 3 min. Wash buffer (1 ml) was added to the lysed cells, and cell nuclei were centrifuged at 500xg for 5 min at 4°C. The nuclei pellet was resuspended in 250 ul of 1x Nuclei buffer. Cell nuclei were then counted using Trypan blue staining. Re-suspended cell nuclei (10-15k) were used for transposition and loaded into the 10x Genomics Chromium Single Cell system. Libraries were amplified with 10 PCR cycles and were sequenced on an Illumina NextSeq with ~200 million reads per library. Sequencing data was processed through the Cell Ranger ATAC 1.1.0 pipeline (10x Genomics) with default parameters.

### Single-cell RNA-seq data preprocessing

Raw scRNA-seq data were processed with the Cell Ranger software (*70*)(version 3.1) for formatting reads, demultiplexing samples, genomic alignment, and generating the cell-by-gene count matrix. First, the ‘cellranger mkfastq’ function was used to generate fastq files from BCL files. Second, the ‘cellranger count’ function was used to process fastq files for each library using default parameters and the mm10 mouse reference index provided by 10x Genomics. Finally, we obtained the cell-by-gene count matrix for each library, and used this for all downstream analysis.

Using the Seurat (*71*) R package, we created Seurat objects for each sample with the cell-by-gene count matrix using the function ‘CreateSeuratObject’ (min.cells = 3, min.features = 200). After visual analysis of the violin plot of the total counts for each cell, we filtered out cells with nCount_RNA < 600 or nCount_RNA > 6000. Next, we calculated the fraction of mitochondrial genes for each cell and filtered out the cells with a mitochondrial fraction > 8%. Finally, we predicted multiplet artifacts and removed potential doublet cells using Scrublet (*72*) for each sample. As a result, 6609 (P8 Ctrl), 6494 (P17 Ctrl), 2607 (P45 Ctrl),12930 (P8 TKO),12413 (P17 TKO), 7531 (P45 KO-replicate 1), and 5886 (P45 KO-replicate 2) were used for downstream analysis.

### Single-cell RNA-seq data analysis

#### Dimensional reduction, clustering and visualization

To integrate the cells from different ages and genotypes, we aligned all cells for each sample and obtained an integrated Seurat object using the Seurat function ‘FindIntegrationAnchors’ and ‘IntegrateData’ with the default parameters, respectively. Using the integrated data, we normalized, log-transformed and scaled the count matrix using the function ‘NormalizeData’ and ‘ScaleData’. We next selected the variable genes using the function ‘FindVariableFeatures’(selection.method = “mvp”) and performed dimension reduction analysis with ‘RunPCA’.

To annotate individual cell types in the integrated dataset, we first clustered all the cells using the function ‘FindNeighbors’ and ‘FindClusters’ with a resolution of 0.3 and 1st-30th dimensions. Then we matched the clusters to major cell types using expression of known cell marker genes. As a result, we identified the following 9 major cell types: Alpha1 tanycytes, Alpha2 tanycytes, Beta1 tanycytes, Beta2 tanycytes, Proliferating tanycytes, Astrocytes, Neurons, Ependymal cells and Oligodendrocyte progenitor cells (OPCs). To visualize the integrated data, we used the 1^st^-30^th^ dimensions to perform nonlinear dimension reduction and obtained UMAP coordinates with the function ‘RunUMAP’.

We further characterized molecularly distinct subtypes of TDNs (tanycytes derived neurons) in Figure 5. First, we restricted our analysis to cells in the neuron cluster and from the following ages and genotypes: P8 Ctrl, P8 TKO, P17 TKO and P45 TKO. P17 Ctrl and P45 Cntrl were excluded from analysis due to the very small numbers of tanycyte-derived neurons present in these datasets. Second, we integrated all the neurons from different conditions using ‘RunHarmony’ in the Harmony R package (*73*). Next, we used the 1 ^st^-10^th^ harmony dimensions to identify neuronal sub-clusters with a resolution of 0.5. Finally, we aggregated the clusters into individual neuronal subtypes based on known neuron markers and RNA velocity results. To visualize tanycyte-derived neurons, we used the 1st-10th harmony dimensions to obtain UMAP coordinates with the function ‘RunUMAP’.

#### Identification of differentially expressed genes

To identify markers for each cell type and differentially expressed genes (DEGs) between Ctrl and TKO samples, we used the Seurat function ‘FindAllMarkers’ and FindMarkers with the options: min.pct = 0.2 or 0.1, logfc.threshold = 0.25. We then retained differential genes with an adjusted p-value of < 0.001.

#### RNA velocity analysis

To characterize cellular differentiation trajectories associated with tanycyte-derived neurogenesis, we used scVelo software (*74*) to perform RNA velocity analysis by comparing levels of spliced and unspliced transcripts. Briefly, we converted bam files for each sample to loom files using a command-line tool (*28*). We then combined these loom files and retained cells which passed filtering in the previous step. Using scVelo, we normalized the spliced and unspliced matrix, filtered the genes and selected the top 1500 variable genes with the function: ‘pp.normalize_per_cell’, ‘pp.filter_genes_dispersion’ and ‘pp.log1p’. Next, we performed a principal component analysis (PCA) and calculated the velocity vectors and velocity graph using the function: ‘pp.moments’(n_pcs=35, n_neighbors=50), ‘tl.recover_dynamics’, ‘tl.velocity’(mode-dynamical’) and ‘tl.velocity_graph’. Finally, we visualized velocities on the previously calculated UMAP coordinates with the ‘pl.velocity_embedding_grid’ function. We applied the same pipeline to analyze RNA velocity in differentiating tanycyte-derived neurons.

#### Cell-cycle stage inference

The function ‘CellCycleScoring’ in the Seurat package was used to calculate cell cycle phase scores (S score and G2/M score), with the G2/M and S phase marker genes obtained from Tirosh et al (*75*).

#### Single-cell trajectory inference

Slingshot (*76*) was applied to infer differentiation trajectories from alpha2 tanycytes to neurons. To construct the trajectory, we included cells in the “Alpha2 tanycytes”, “Proliferating tanycytes” and “Neuron” clusters. We then then ran Slingshot using the dimensionality reduction results (UMAP) identified previously. We set the “Alpha2 tanycytes” cluster as the initial cluster to identify lineages with the function “getLineages” and “getCurves” with default parameters. Finally, we assigned cells to the lineages and calculated pseudo-time values for each cell using the function “slingPseudotime.”

Monocle 2 (*77*) was applied to identify developmentally dynamic genes which are significantly altered along the trajectory. First, we converted the expression matrix to Monocle datasets with the function ‘newCellDataSet’, then we processed and normalized the Monocle datasets following the Monocle recommended pipeline, and finally identified DEGs using the “differentialGeneTest” function with the following criteria: q-value < 1e-10 and expressed cell number > 200.

#### Comparison between tanycyte-derived neurons and mature hypothalamic neurons

To further explore the biological similarity between tanycyte-derived neurons and the broader population of neurons in mouse hypothalamus, we first used the scRNA-seq datasets for mature neurons in hypothalamic arcuate nucleus provided by Campbell, et al (*23*), and downloaded the cell-by-gene matrix and the annotation file of the mature neuronal cell types from the GEO database under the accession GSE93374. The LIGER (*33*) package was used to integrate our tanycyte-derived cells identified in the previous rounds of analysis with these mature hypothalamic neurons, using the default pipeline recommended in the LIGER guidelines (https://macoskolab.github.io/liger/). After LIGER integration, we re-clustered the integrated datasets and calculated new UMAP coordinates using the function ‘FindNeighbors’, ‘FindClusters’ and ‘RunUMAP’ with the following parameters: reduction = “iNMF”, dims = 1:30. Finally, we made an alluvial plot to visualize the alignments between the tanycyte-derived neuronal sub-types (6 subtypes), LIGER clusters (14 clusters) and the mature arcuate neuronal cell types (34 subtypes).

#### Single-cell ATAC-seq data preprocessing

We processed sequencing output data using the Cell Ranger ATAC software (v.1.0) for alignment, de-duplication, and identification of transposase cut sites. First, the ‘cellranger-atac mkfastq’ function was used for generating fastq files from BCL files. Second, the ‘cellranger-atac count’ function was used to process the fastq files for each library using default parameters and the mouse mm10 reference index provided by 10x Genomics (refdata-cellranger-atac-GRCh38-1.2.0). Finally, we obtained the barcoded, aligned, and Tn5-corrected fragment file (fragments.tsv.gz) for each library and used these for downstream analysis.

### Single-cell ATAC-seq data analysis

#### Generating union peaks

We generated the cell-by-peaks matrix for each sample using the same method as described in Satpathy, A. T. et al (*70*). First, we constructed 2.5-kb tiled windows across the mm10 genome using the local script. Next, a cell-by-window sparse matrix was computed by counting the Tn5 insertion overlaps for each cell, and this matrix was then binarized and inputted to Signac package (0.2.5) to create a Seurat object using ‘CreateSeuratObject.’ Second, we normalized the cell-by-window matrix by TF-IDF methods using ‘RunTFIDF’ and ran a singular value decomposition (SVD) on the TF-IDF normalized matrix with ‘RunSVD.’ We retained the 2^nd^ to 30^th^ dimensions, and identified clusters using SNN graph clustering with ‘FindClusters’ with a resolution of 0.3. Third, to identify high-quality peaks for each cluster in each sample, we called peaks for each cluster using MACS2 (*78*) with the command: ‘-shift −75 --extsize 150 -- nomodel --callsummits --nolambda --keep-dup all -q 0.05’. The peak summits were then extended to 250 bp on either side to a final width of 500 bp and then filtered by the mm10 v2 blacklist regions (https://github.com/Boyle-Lab/Blacklist/blob/master/lists/mm10blacklist.v2.bed.gz). The top 50,000 peaks for each cluster in each sample were kept, converted to Granges, and merged into a union peak set with the function ‘reduce’ in the GenomicRanges package. Finally, we obtained 107,377 union peaks, and generated a cell-by-peak sparse matrix for all these cells for downstream analysis.

#### Filter cells by TSS enrichment, unique fragments, and nucleosome banding

We calculated the TSS enrichment, unique fragments, and nucleosome banding for each cell using the Signac package. The cell-by-peak sparse matrices were inputted to the ‘CreateSeuratObject’ function to create a Seurat object with default parameters. Then we filtered the cells using the following criteria: 1) The number of unique nuclear fragments > 1000; 2) TSS enrichment score > 2; 3) nucleosome banding score < 4; 4) blacklist_ratio < 0.05. As a result, 8948 (P8 Ctrl) and 13337 (P8 TKO) cells were identified and used for downstream analysis.

#### Dimensional reduction, clustering and visualization

The Harmony package was applied to integrate the scATAC-seq data from control and *Nfia/b/x* TKO samples. First, we put the Seurat object created in the previous step into the Signac process pipeline. We normalized and obtained a low-dimensional representation of the cell-by-peak matrix using the functions ‘FindTopFeatures’, ‘RunTFIDF’ and ‘RunSVD’. Next, we integrated all the cells from both genotypes (control and TKO) using the ‘RunHarmony’ function with the options: dim.use = 2:50, group.by.vars = ‘condition’, reduction = ‘lsi’ and project.dim = FALSE. Third, we used the 2^nd^-30^th^ harmony dimensions to identify clusters with a resolution of 0.8, and used the same harmony dimensions to calculate the UMAP coordinates for visualization.

To annotate the cell types for each cluster, we used the integration method in the Seurat package to transfer the previously annotated cell-type labels from scRNA-seq data to scATAC-seq data. First, we estimated RNA-seq levels using the function ‘CreateGeneActivityMatrix’ from the scATAC-seq data using the mm10 genome build gtf file. Next, we found anchors between the scATAC-seq datasets (P8 Ctrl and P8 TKO) and the corresponding scRNA-seq datasets (P8 Ctrl and P8 TKO) using the function ‘transfer.anchors.’ Then with the ‘TransferData’ function, we obtained a matrix of celltype predictions and prediction scores for each cell in the scATAC-seq dataset. We further filtered the cells with max(prediction score) < 0.5. Finally, for each cluster, we calculated the number of cells for each predicted cell type, and set its final annotation based on the cell type that was most highly represented in the cluster. Using this approach, we identified the following 9 major cell types: Alpha1 tanycytes, Alpha2 tanycytes, Beta1 tanycytes, Beta2 tanycytes, Proliferating tanycytes, Astrocytes, Neurons, Ependymal cells and OPCs.

#### Global NFI activity and footprint analysis

We inferred global NFI activity with the chromVAR R package (*79*). The raw cell-by-peak matrix from the total cells was used as input to chromVAR. The mm10 reference genome was used to correct GC bias. We used the mouse NFI Motifs (including Nfia, Nfib and Nfix) in the TransFac2018 database to generate the transcription factor (TF) z-score matrix. The z-score for each cell was used to visualize the global NFI activity using the previously calculated UMAP coordinates.

To analyze the NFI footprint in alpha2 tanycytes, we used the same methods described in Corces et al. (*80*). First, we used the NFI motifs and all accessible regions to predict the NFI binding sites with the function ‘match motifs’ in the motif matching R package. Second, we calculated the Tn5 insertion bias around every NFI binding site. We generated the aggregated observed 6-bp hexamer table relative to the ± 250 bp region from all motif centers, and we also calculated the aggregated expected 6-bp hexamer table from the mm10 genome. We then obtained the observed/expected (O/E) 6-bp hexamer table by dividing these two hexamer tables. Finally, we calculated the observed Tn5 insertion signal at ± 250 bp from the motif center, and normalized the signal using the O/E 6-bp hexamer table to obtain the final Tn5 bias-corrected signal.

#### Differential peak analysis

To explore which ATAC regions are changed following Nfia/b/x loss of function, we applied the MAnorm algorithm (*81*) to perform differential peak analysis between control and *Nfia/b/x* TKO alpha2 tanycytes. First, we selected cells in the ‘Alpha2 tanycytes’ cluster and then separated these cells by genotype (control and *Nfia/b/x* TKO). Second, we aggregated the cells of the same genotype by summing the count signals for each peak, then created a new condition-by-peak count matrix, and put it into the MAnorm pipeline. Finally, we performed the MAnorm test and identified differential peaks (4563 peaks enriched in Ctrl and 1333 peaks enriched in TKO) with the cutoff: LOG_P > 25, and abs(M_value_rescaled) > 0.5.

#### De novo motif enrichment analysis

HOMER software (*82*) was applied to identify motifs enriched in the differential ATAC regions between control and *Nfia/b/x* TKO alpha2 tanycytes. We analyzed the up-regulated peaks and down-regulated peaks separately using the Homer function ‘findMotifsGenome.pl’ with the default options, except: mm10, -size given, -mask.

#### Identification of genes directly regulated by Nfia/b/x

To further explore the biological function of NFI factors in alpha2 tanycytes, we developed a method to infer potential Nfia/b/x targets with the information in scRNA-seq and scATAC-seq data. Our methods included the following 3 steps:

##### 1. Identification of Nfia/b/x-binding regions

In the previous motif enrichment analysis, we found that NFI motifs are enriched in the down-regulated peaks (*Nfia/b/x* TKO/control), so in the first step, we aimed to identify which down-regulated peaks are bound by NFI factors. Using the NFI motif information in the TRANSFAC2018 database, we first scanned the NFI motifs in the 4563 down-regulated peaks with the function ‘matchMotifs’. Next, with the Tn5 insertion signal in P8 Ctrl alpha2 tanycytes, we calculated the footprint occupancy score (FOS) (*83*) for each predicted NFI binding region, and filtered out the regions with FOS < 2. Finally, we kept only the NFI-binding peaks which contain NFI binding sites, and used them for downstream analysis.

##### 2. Identification of promoters associated with Nfia/b/x binding regions

To identify genes that are potentially regulated by these NFI binding regions, we used the Cicero algorithm (*84*) to identify all the distal elements–promoter connections genome-wide. First, we converted the cell-by-peak sparse binary matrix into the Cicero pipeline with the functions ‘make_atac_cds’, ‘detectedGenes’ and ‘estimatedSizeFactors’. Next, we created low-overlapping cell groups based on the KNN nearest-neighbors in the UMAP dimension, and aggregated signals for each cell group with the function ‘make_cicero_cds’. We then calculated the correlation between each peak–peak pair using the function ‘run_cicero’ with default parameters. Third, we annotated the peak pairs using ‘annotate_cds_by_site’ with mm10 gtf files. We kept the peak pairs with the following criteria: 1) one of the peaks overlapped with ±2 kb of TSS region; and 2) one of the peaks contained at least one NFI binding motif. Finally, we identified NFI-related distal elements–promoter connections from the peak pairs if their co-accessibility score >0.03, or <-0.03 and their distance < 150kb.

##### 3. Inference of potential Nfia/b/x targets by integrating with scRNA-Seq data

In this step, we aimed to integrate the NFI-related distal elements–promoter connections and differential genes following loss of function of *Nfia/b/x* to identify NFI target genes. First, we selected enhancer-promoter pairs from the distal elements– promoter connections in Step 2 with co-accessibility scores > 0.03. If the gene associated with the promoter in question was down-regulated following loss of function of *Nfia/b/x*, we treated these genes as potential Nfia/b/x targets.

Conversely, we selected silencer-promoter pairs with co-accessibility scores < −0.03. If the promoter genes were up-regulated following loss of function of *Nfia/b/x*, we also treated these genes as potential Nfia/b/x targets. Using this approach, we identified 63 NFI target genes.

### GO term analysis

To understand the biological functions associated with genes dynamically expressed during the process of alpha2 tanycyte-derived neurogenesis, we applied GOrilla algorithm (*85*) to identify enriched Gene Ontology terms for each gene cluster using the default parameters (P-value threshold = 0.001, ontology = ‘Process’). The output of Gene Ontology terms from GOrilla were further processed by REVIGO (*86*) to remove redundant terms. This pipeline was also used to identify the GO term enrichment in NFI-regulated gene sets.

### Brain slice preparation and cell type identification

To investigate the electrophysiological characteristics of tanycyte-derived and other hypothalamic neurons, acute brain slices were generated as previously described (*37*). *Rax-CreER;Nfia^lox/lox^;Nfib^lox/lox^;Nfix^lox/lox^;CAG-lsl-Sun1-GFP* mice (P15-P97, male) were anesthetized with isoflurane, decapitated, and the brains were rapidly removed and chilled in ice-cold sucrose solution containing (in mM): 76 NaCl, 25 NaHCO_3_, 25 glucose, 75 sucrose, 2.5 KCl, 1.25 NaH_2_PO_4_, 0.5 CaCl_2_, and 7 MgSO_4_; pH 7.3. Acute brain slices (300 μm) including the hypothalamus were prepared using a vibratome (VT-1200s, Leica) and transferred to warm (32-35°C) sucrose solution for 30 minutes for recovery. The slices were transferred to warm (32-34°C) artificial cerebrospinal fluid (aCSF) composed of (in mM): 125 NaCl, 26 NaHCO_3_, 2.5 KCl, 1.25 NaH_2_PO_4_, 1 MgSO_4_, 20 glucose, 2 CaCl_2_, 0.4 ascorbic acid, 2 pyruvic acid, and 4 L-(+)-lactic acid; pH 7.3, 315 mOsm, and allowed to cool to room temperature (RT). All solutions were continuously bubbled with 95% O_2_/5% CO_2_. For whole-cell patch-clamp recordings, slices were transferred to a submersion chamber on an upright microscope (Zeiss AxioExaminer, Objective lens: 5x, 0.16 NA and 40x, 1.0 NA) fitted for infrared differential interference contrast (IR-DIC) and fluorescence microscopy. Slices were continuously superfused (2-4 ml/min) with warm, oxygenated aCSF (32-34°C). Hypothalamic areas and cells were identified under a digital camera (Sensicam QE; Cooke) using either transmitted light or green fluorescence. Tanycytes were identified as GFP-positive cells located in the ependymal cell layer at the 3^rd^ ventricle. Tanycyte-derived cells were identified as GFP-positive cells located in the hypothalamic parenchyma but not in the ependymal cell layer. GFP-negative hypothalamic neurons in the hypothalamic parenchyma, among which were intermingled the sparse tanycyte-derived cells, were targeted as control neurons.

### Whole-cell recordings and analysis

Borosilicate glass pipettes (2-4 MΩ) were filled with an internal solution containing (in mM): 2.7 KCl, 120 KMeSO_4_, 9 HEPES, 0.18 EGTA, 4 MgATP, 0.3 NaGTP, 20 phosphocreatine(Na), pH 7.3, 295 mOsm. Biocytin (0.25% weight/volume) was added to the internal solution for post-hoc morphological characterization. Whole-cell patchclamp recordings were conducted through a Multiclamp 700B amplifier (Molecular Devices) and an ITC-18 (Instrutech) which were controlled by customized routines written in Igor Pro (Wavemetrics). The series resistance averaged 14.2 ± 5.8 MΩ SD (*n* = 81 cells, 12 mice, all < 36 MΩ, no significant difference between cell types or age groups, p > 0.05, Mann-Whitney U test), and was not compensated. The input resistance was determined by measuring the voltage change in response to a 1 s-long - 100 pA hyperpolarizing current step. The current-spike frequency relationship was measured with a series of depolarizing current steps (1 s-long, 0-50 pA, 10 pA increments, 5 s interstimulus intervals). For each current intensity, the total number of action potentials exceeding 0 mV generated during each step was measured and then averaged across the three trials. Spontaneous postsynaptic currents (sPSCs) were measured in voltage-clamp mode at −70 mV. sPSCs were recorded for 25 sec (250 ms-long current traces, 100 times), and ~110 events, on average, were recorded per cell. High amplitude, high frequency depolarizing current steps (10 nA at 100 Hz for 100 ms) were injected into the recorded cells at the end of recording, to increase efficiency of biocytin infusion (*87*). All signals were low-pass filtered at 10 kHz and sampled at 20 kHz for voltage traces and 100 kHz for series resistance and sPSCs measurements.

### Electrophysiology data analysis and statistical testing

Data analysis was performed in Igor Pro (WaveMetrics), Excel (Microsoft), ImageJ (NIH), and Minhee analysis (https://github.com/parkgilbong/Minhee_Analysis_Pack). Data are presented as the mean ± SEM unless otherwise noted. A Mann-Whitney U test was used to compare membrane properties and sPSCs frequencies between cell types and between age groups. Spearman’s Rho test was used to determine the correlation between sPSC frequency and cell location. The location of the cells (distance to tanycytic layer) was measured from low (5x, 0.16 NA) and high (40x, 1.0 NA) magnification images of the recorded cells using ImageJ. The statistical difference in current-spike frequency relationships was tested by using a two-way ANOVA test with Bonferroni correction. The sPSC events were automatically detected by Mini analysis software with a 10 pA amplitude threshold. In the figures, the statistical significance is expressed as follows: *p < 0.05; **p < 0.01; or ***p < 0.001.

### Visualization of biocytin-filled cells

Following the electrophysiological experiments, slices were fixed in 4% PFA in 0.01 M PBS at least overnight. After rinsing with PBS, slices were incubated in 0.01 M PBS blocking solution containing 2% Triton X-100 (Sigma-Aldrich) and 5% normal donkey serum (NDS) for 1 h at RT. To visualize biocytin-filled cells, slices were next incubated with a blocking solution containing 1% Triton X-100, 5% NDS, chicken anti-GFP antibody (1:1,000, Aves, Cat. No. GFP-1020), and AlexaFluor 647-conjugated streptavidin overnight on shaker at 4°C. The following day, slices were rinsed with 0.01 M PBS, and incubated with AlexaFluor 488-conjugated donkey-anti-chicken (1:500, Jackson ImmunoResearch, Cat. No. 705-745-155) for 2 h at RT. After rinsing, slices were mounted with Aqua-Poly/Mount (18606-20, Polysciences). A subset of slices was co-stained with mouse anti-NeuN (1:300, Millipore, MAB377) and AlexaFluor 568-conjugated donkey anti-mouse (1:300, ThermoFisher, A10037) antibodies to confirm neuronal identity of the biocytin-filled cells. Fluorescence images were taken with a confocal microscope (LSM 800, Zeiss; 20x objective lens) as z-stack (2 μm-interval) tiled images to cover the extent of the cell’s dendritic and axonal processes.

---

**Supplementary Figure 1:**
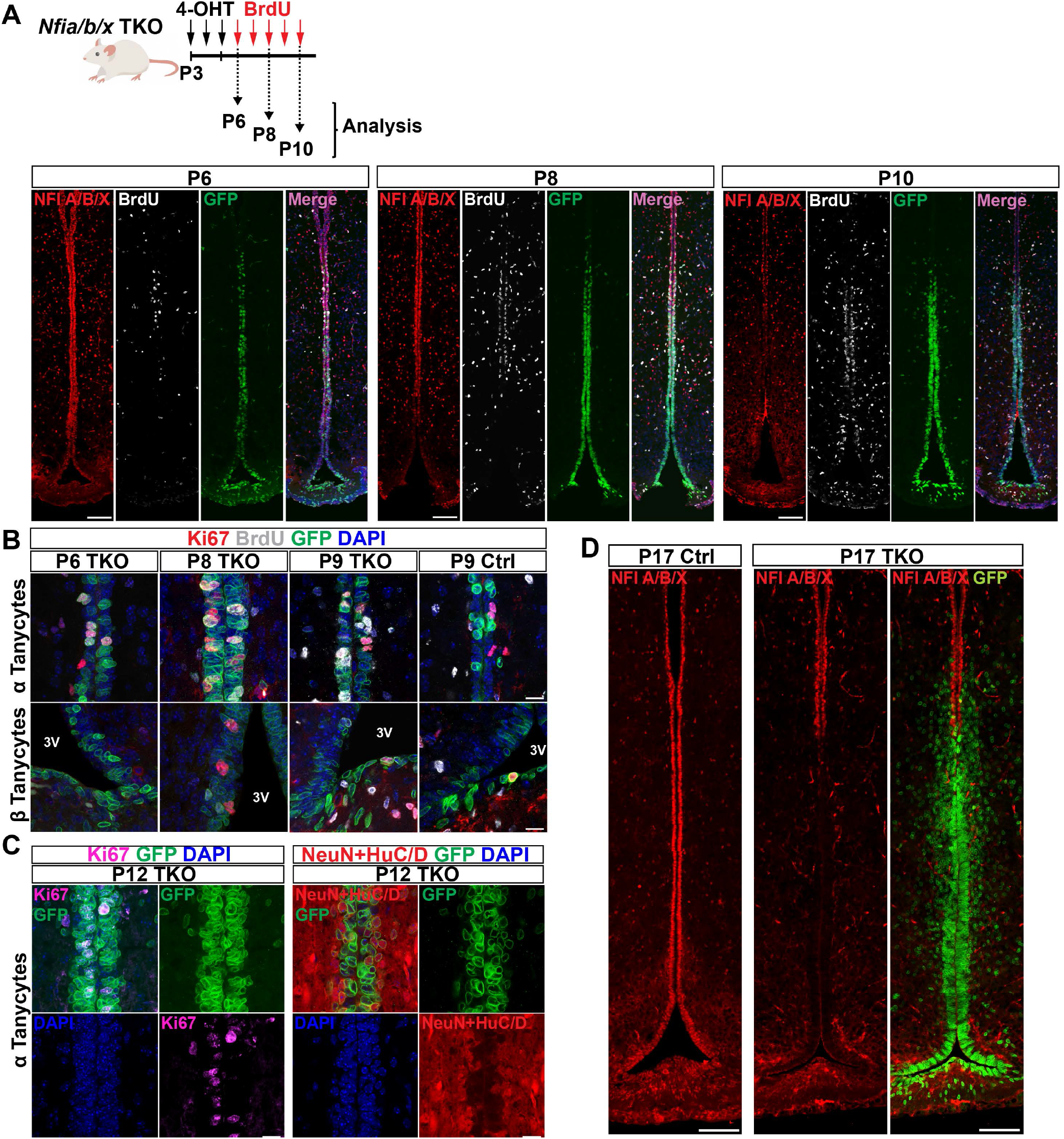
Time course of tamoxifen-dependent loss of Nfia/b/x protein expression and induction of proliferation in *Nfia/b/x* mutant mice. A. Loss of Nfia/b/x immunoreactivity and induction of BrdU labeling following tamoxifen treatment of *Rax-CreER;Nfia^lox/lox^;Nfib^lox/lox^;Nfix^lox/lox^;CAG-lsl-Sun1-GFP* mice at P6, P8 and P10. B. Induction of proliferation occurs in alpha tanycytes before being detectable in beta tanycytes. C. The alpha tanycytic ventricular zone thickens at P12 and Ki67+ cells are localized in the most superficial layer while immunoreactivity to HuC/D and/or NeuN is detected at the layer closest to the parenchyma. Antibodies to HuC/D and NeuN were combined for this analysis. D. Near-complete, tanycyte-specific loss of Nfia/b/x immunoreactivity by P17. Scale bars: A, D=100 μm; B,C=20 μm.

**Supplementary Figure 2:**
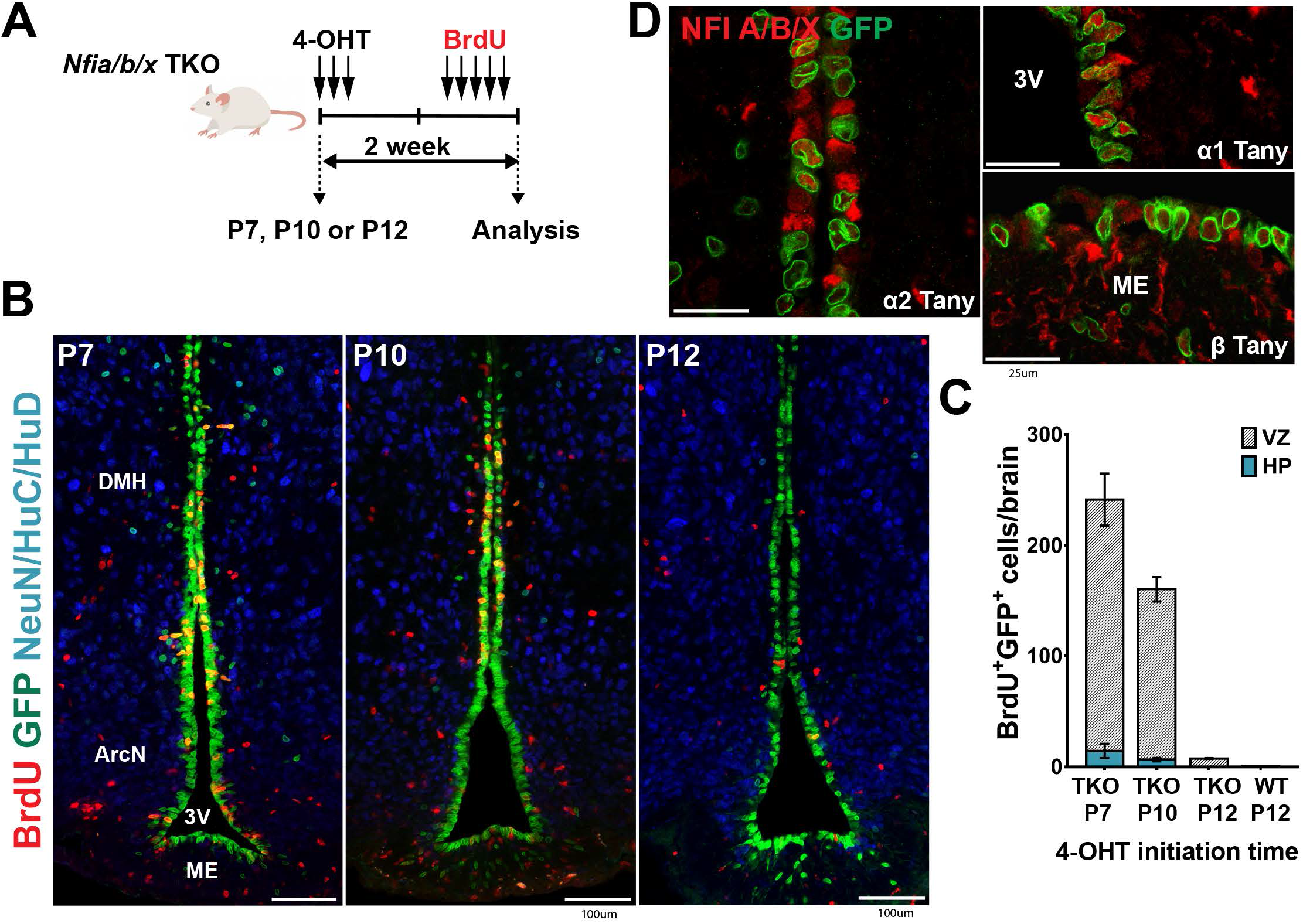
Tamoxifen-dependent Cre activation does not effectively induce proliferation in *Nfia/b/x* mutant mice at P12. A. Schematic for tamoxifen-dependent disruption of Nfia/b/x function in *Rax-CreER;Nfia^lox/lox^;Nfib^lox/lox^;Nfix^lox/lox^;CAG-lsl-Sun1-GFP* mice at P7, P10, and P12. B. Induction of proliferation and neurogenesis by *Nfia/b/x* deletion at P7, P10, and P12. C. Mosaic loss of Nfia/b/x immunostaining following Cre activation at P12. D. BrdU labeling in ventricular zone (VZ) and hypothalamic parenchyma (HP) following tamoxifen treatment at P7 (n=3 mice), P10 (n=4 mice) and P12 (n=2-3 mice). Scale bars: B=100 μm, D=25 μm. Abbreviations: 3V=third ventricle, DMH=dorsomedial hypothalamus, ME=median eminence.

**Supplementary Figure 3:**
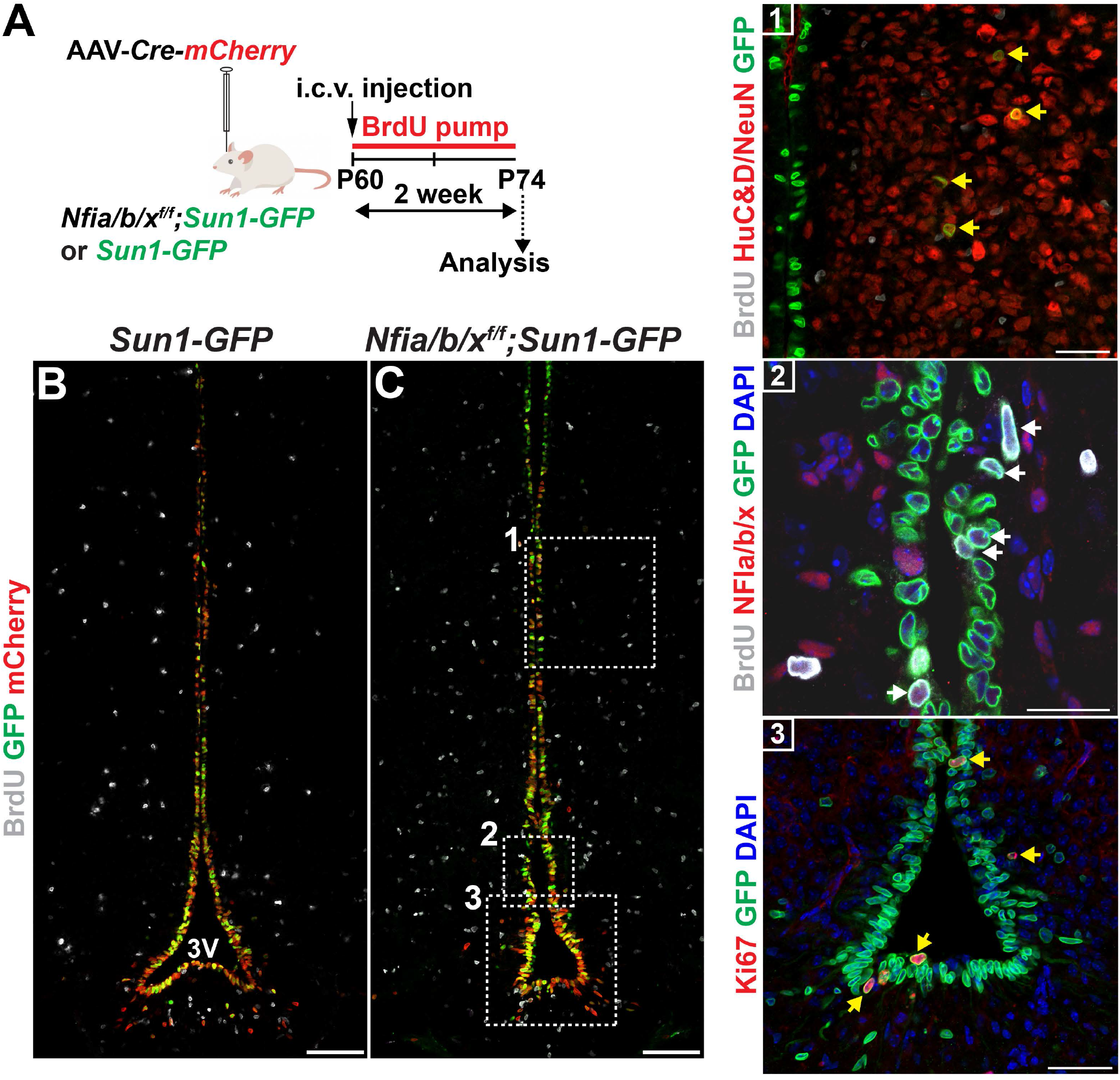
AAV-Cre-mediated deletion of NFI genes induced tanycyte proliferation in adult mice. A. Cre-dependent AAV-based deletion strategy. AAV1-Cre-mCherry was injected into the lateral ventricles of *Nfia^lox/lox^;Nfib^lox/lox^;Nfix^lox/lox^;CAG-lsl-Sun1-GFP (Nfia/b/xf/f;Sun1-GFP*) or *CAG-lsl-Sun1-GFP (Sun1-GFP*, as controls) mice at P60, followed by BrdU infusion over 2 weeks using an osmotic minipump. B-C. Confirmation of viral infection with immunolabeling for mCherry and GFP induction in the ventricular layer of both controls (B) and NFI knockout mice (C). A subset of parenchymal GFP+ cells were immunoreactive for HuC/D and NeuN (yellow arrows in inset 1), but BrdU-negative. Only NFI knockout mice show BrdU and GFP double-positive cells in the tanycytic layer (white arrows in inset 2). Induction of proliferation was confirmed with co-immunostaining for Ki67 and GFP in the tanycytic layer (yellow arrows in inset 3). Scale bars: B, C=100 μm; inset 1, 3=50 μm; inset 2=25 μm.

**Supplementary Figure 4:**
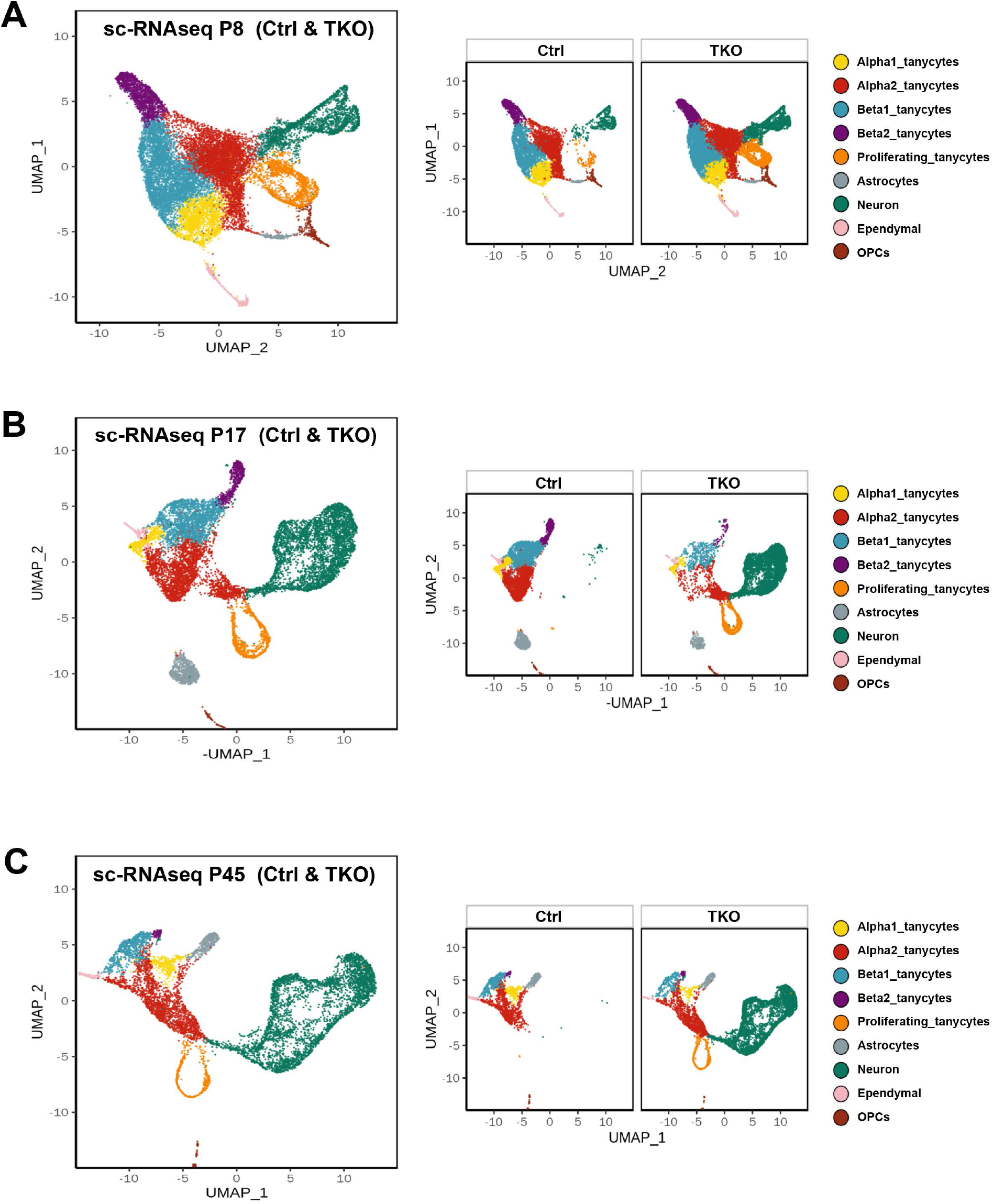
Clustering of GFP-positive cells at different ages. P8 (A), P17 (B), P45 (C).

**Supplementary Figure 5:**
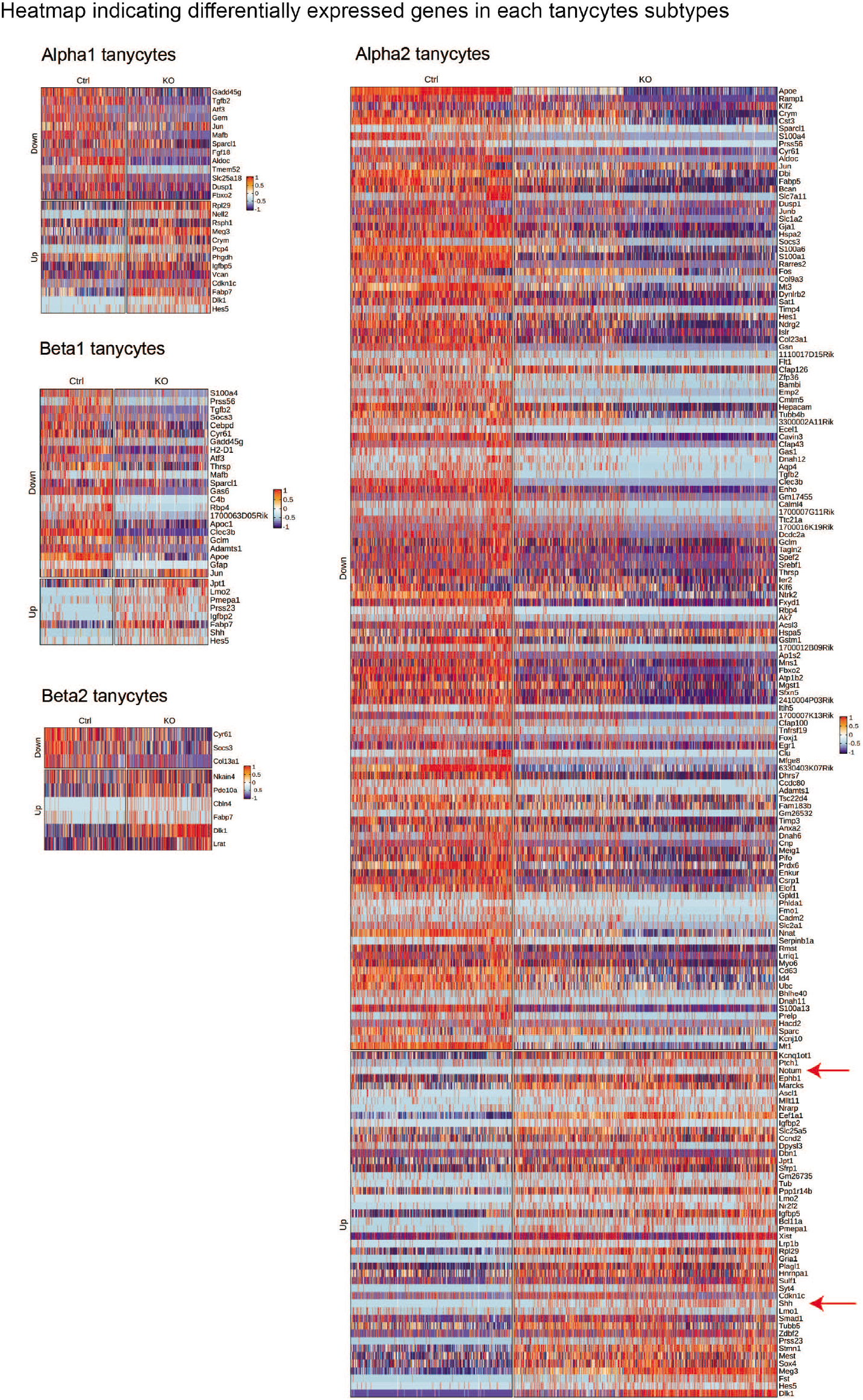
Heatmap showing differentially expressed genes in each tanycytes subtype. Major changes in gene expression observed in TKO alpha2 tanycytes. Up-regulated *Notum* and *Shh* expression in alpha2 tanycytes is highlighted (Red arrows).

**Supplementary Figure 6:**
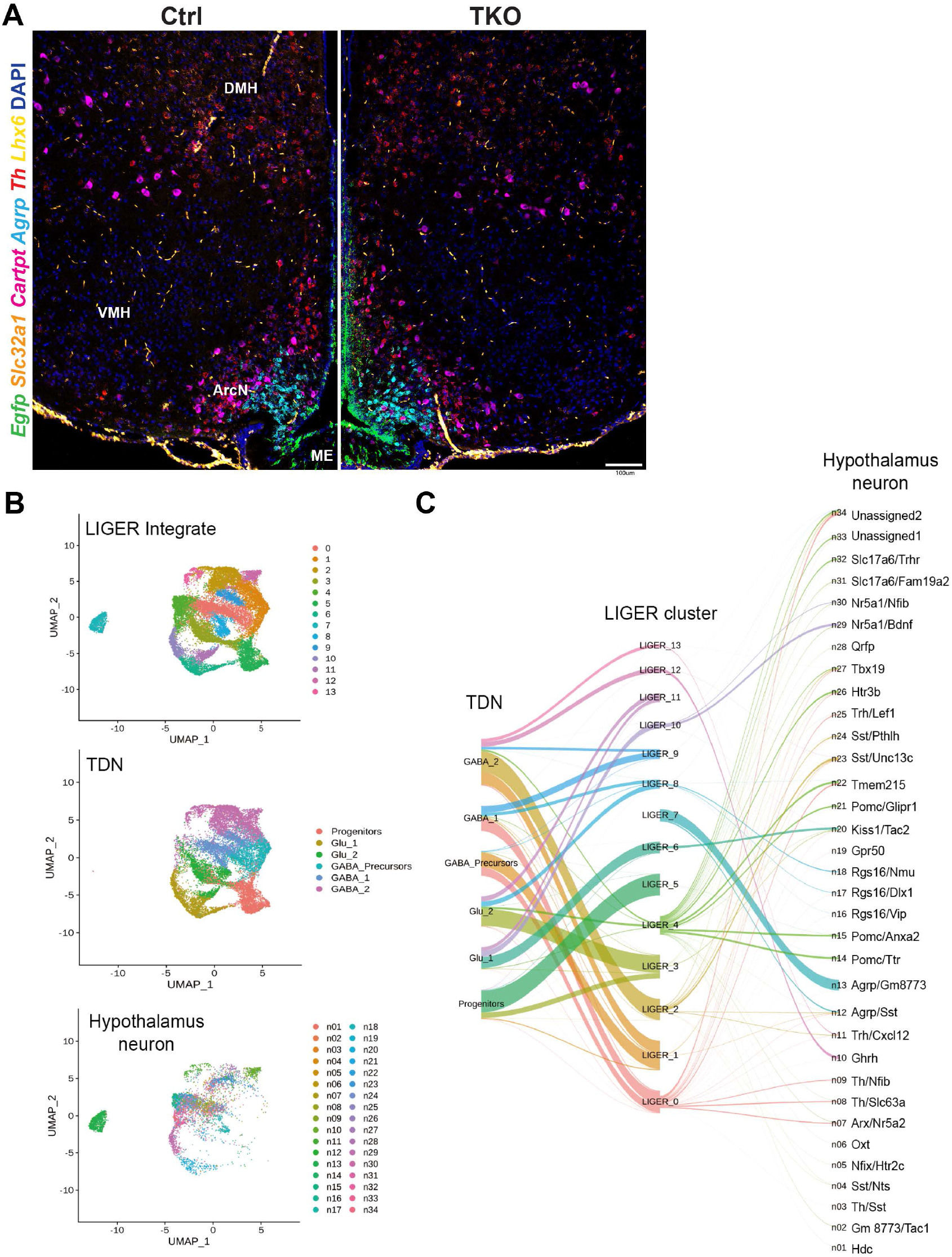
Hypothalamic neuronal distribution and comparison analysis with a published scRNA-Seq data from ArcN neurons. A. No obvious change is observed in the overall distribution of hypothalamic neurons in *Nfia/b/x*-deficient mice by Hiplex RNAscope analysis. B. LIGER-based comparison of P8, P17 and P45 tanycyte-derived neurons with scRNA-Seq data from normal and low fat chow-fed mice from ArcN (Campbell, et al. 2017), with tanycyte-derived neurons and ArcN neurons plotted separately. C. Alluvial plot indicating relationships between identified clusters of tanycyte-derived neurons (this study) and ArcN neurons (from Campbell, et al. 2017). Scale bars: A=100 μm.

**Supplementary Figure 7:**
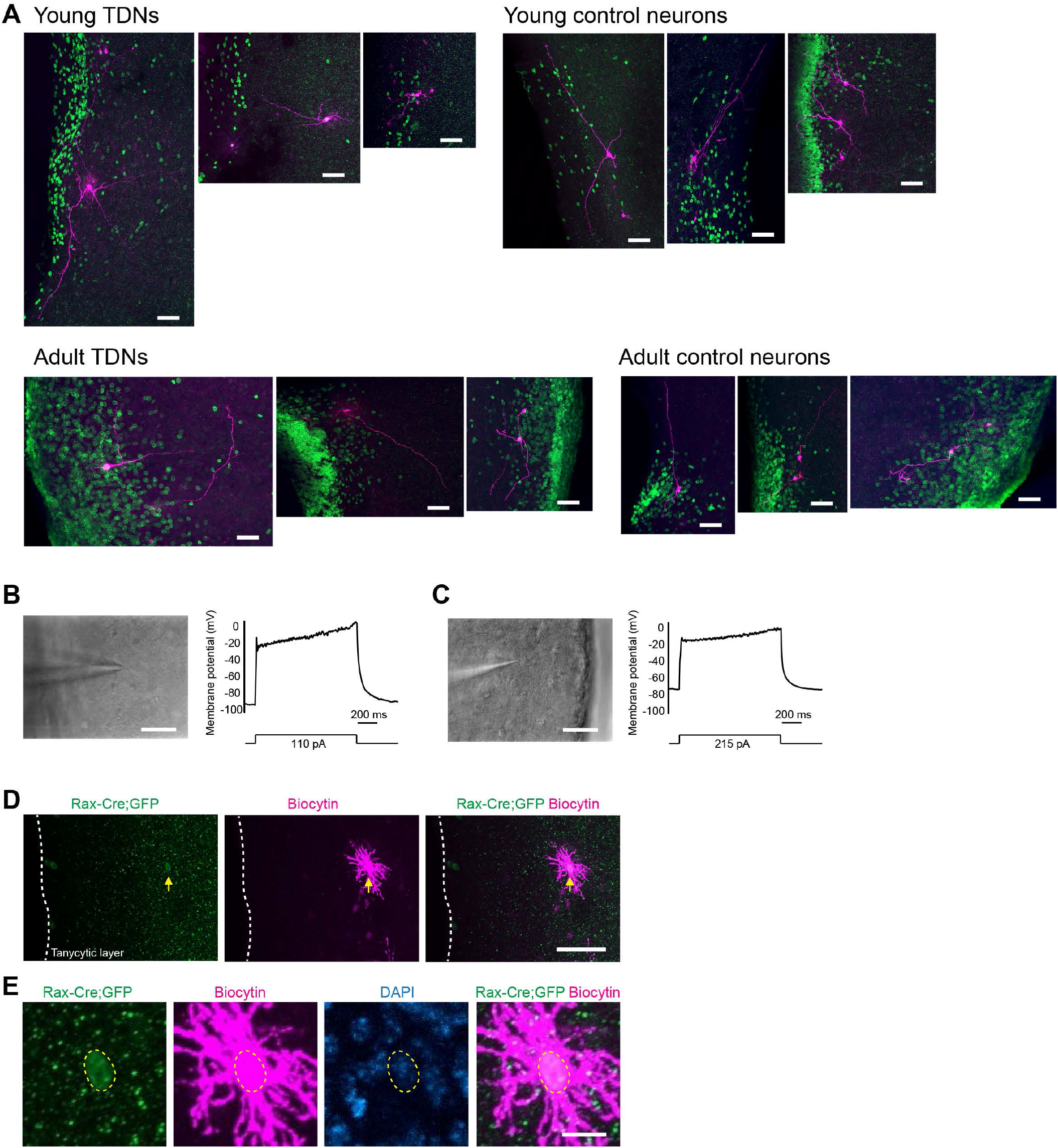
Morphology of biocytin-filled tanycyte-derived and control neurons and non-neuronal tanycyte-derived cells. A. Three examples of biocytin-filled tanycyte-derived and control neurons (magenta). GFP+ tanycyte-derived neurons and GFP-control neurons were recorded from TKO brain slices. B.C. Two tanycyte-derived cells in the hypothalamic parenchyma (left images, DIC images) of young *NfIa/b/x*-deficient mice (B: P18 mouse, C: P16 mouse) that did not fire action potentials in response to step depolarizations of current (right). Low (D) and high (E) magnification confocal images of the cell shown in **C** visualized by biocytin-streptavidin staining. This cell expressed GFP but showed glial cell-like morphology. Scale bars: A-D=50 μm, E=10 μm.

**Supplementary Figure 8:**
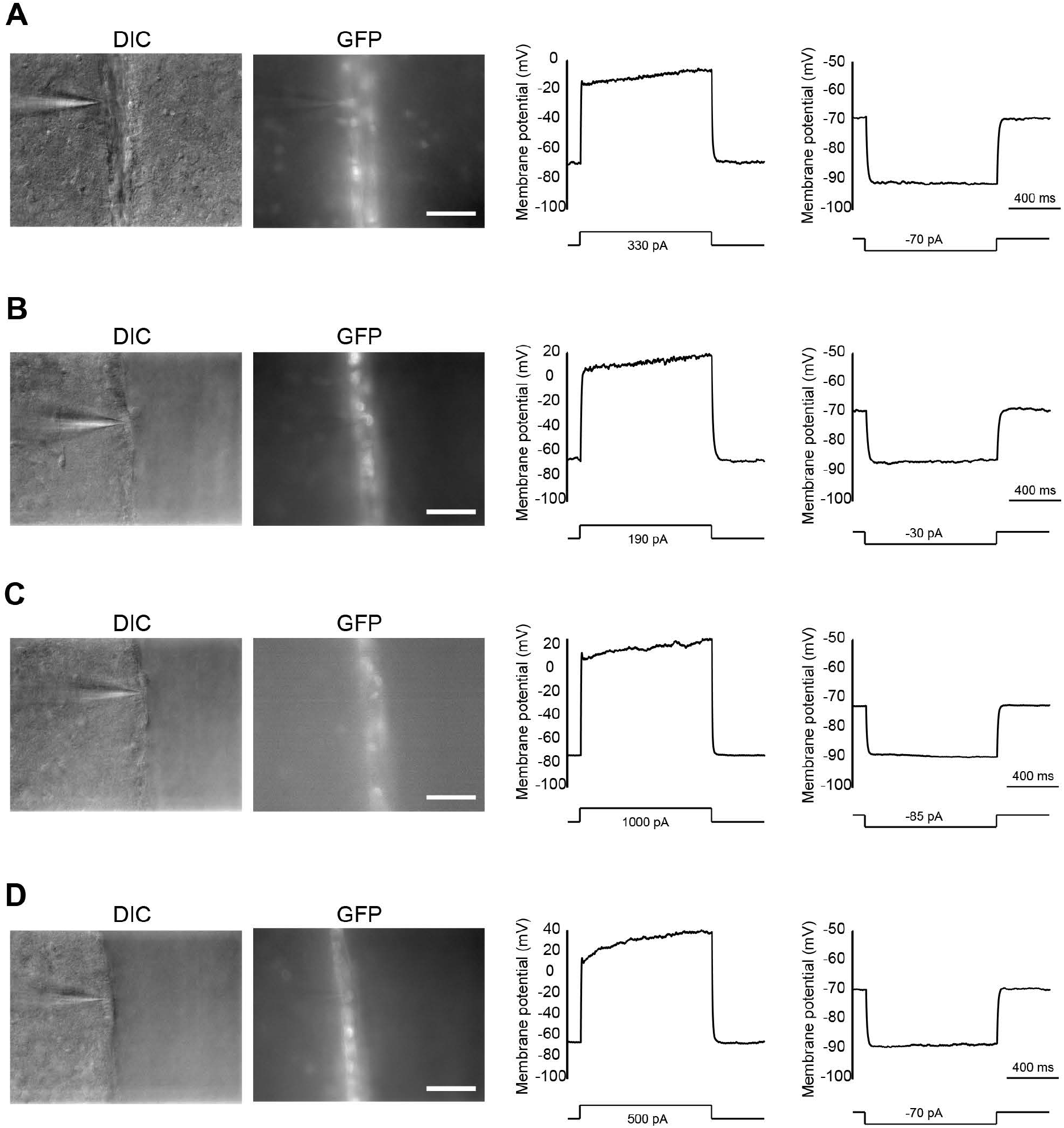
Tanycytes in the tanycytic layer are non-spiking cells in *NFIa/b/x* TKO mice. A-D. DIC and GFP fluorescence images of four cells located in the tanycytic layer and their membrane potential responses to depolarizing and hyperpolarizing current steps. Cells were recorded from P18 (**A**) and P42 (**B-D**) *NF/a/b/x* TKO mice. Note that these cells did not fire action potentials to depolarizing current steps. Scale bars: A-D=50 μm

**Supplementary Figure 9:**
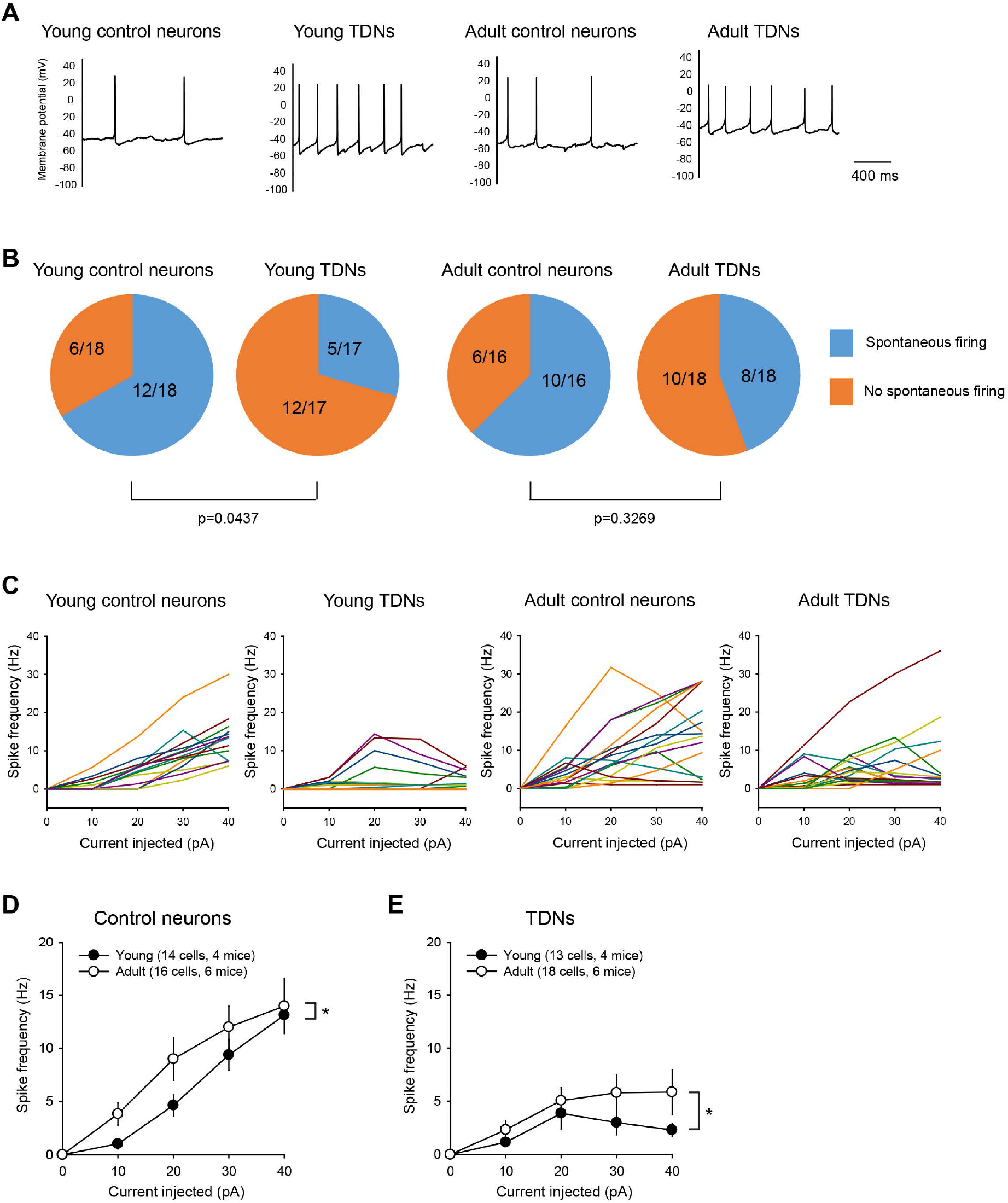
Tanycyte-derived neurons fire spontaneous action potentials and show increased firing with age. A. Examples of recorded cells showing spontaneous action potentials (sAPs) at their resting potentials. B. Proportion of neurons displaying sAPs is higher in control neurons than tanycyte-derived neurons in young mice (p = 0.0437, Fisher’s exact test), whereas there is no difference in adult mice (p = 0.3269, Fisher’s exact test). C. Current-spike frequency relationship plots for individual young control neurons (14 cells from 4 mice), young tanycyte-derived neurons (13 cells from 4 mice), adult control neurons (16 cells from 6 mice), and adult tanycyte-derived neurons (18 cells from 6 mice). The summary data is shown in Figure 6G. One neuron (plotted in bold orange) did not fire action potentials in response to 10-40 pA current injections, but generated action potentials following a 340 pA current injection and was therefore categorized as a tanycyte-derived neuron. D.,E. Current-spike frequency relationship plots showing differences in spike frequency of control neurons (D) or tanycyte-derived neurons (E) between young and adult mice. Both control neurons (p = 0.0304, Two-way ANOVA) and tanycyte-derived neurons (p = 0.02977, Two-way ANOVA) showed significantly higher spike frequency in cells from adult mice compared to cells from young mice.

## Supplemental Tables

**Table S1**: Antibodies used for IHC.

**Table S2**: Number of each cell type identified in each scRNA-Seq library.

**Table S3**: Marker genes in controls and *Nfia/b/x*-deficient tanycytes at P8, P17, and P45.

**Table S4**: Differentially expressed genes in *Nfia/b/x*-deficient alpha2 tanycytes.

**Table S5**: Differentially expressed genes in six clusters separated across pseudotime between alpha2 tanycytes and neuronal precursors.

**Table S6**: HOMER analysis showing most enriched motifs in up/down peaks.

**Table S7**: Differentially expressed genes in *Nfia/b/x*-deficient alpha2 tanycytes at P8.

**Table S8**: Differentially accessible regions in *Nfia/b/x*-deficient alpha2 tanycytes at P8.

**Table S9**: NFI-target genes differentially accessible in *Nfia/b/x*-deficient alpha2 tanycytes.

**Table S10**: Numbers of cells in each cluster of tanycyte-derived neurons and number of Lepr-positive cells in each cluster.

**Table S11**: Marker genes for each cluster in tanycyte-derived neurons.

**Table S12**: Probes used for HiPlex smfISH.

**Table S13**: Genes enriched in LIGER clusters generated by comparing scRNA-Seq profiles of tanycyte-derived neurons and scRNA-Seq data of ArcN neurons.

**Table S14**: Electrophysiological properties of control GFP-negative neurons and tanycyte-derived neurons recorded from young (P15-P19) and adult (P86-P97) NFI TKO mice.

## References

1. E. M. Rodríguez, J. L. Blázquez, F. E. Pastor, B. Peláez, P. Peña, B. Peruzzo, P. Amat, Hypothalamic tanycytes: a key component of brain-endocrine interaction. Int. Rev. Cytol. 247, 89–164 (2005).

2. M. Bolborea, N. Dale, Hypothalamic tanycytes: potential roles in the control of feeding and energy balance. Trends Neurosci. 36, 91–100 (2013).

3. D. A. Lee, J. L. Bedont, T. Pak, H. Wang, J. Song, A. Miranda-Angulo, V. Takiar, V. Charubhumi, F. Balordi, H. Takebayashi, S. Aja, E. Ford, G. Fishell, S. Blackshaw, Tanycytes of the hypothalamic median eminence form a diet-responsive neurogenic niche. Nat. Neurosci. 15, 700–702 (2012).

4. S. C. Robins, I. Stewart, D. E. McNay, V. Taylor, C. Giachino, M. Goetz, J. Ninkovic, N. Briancon, E. Maratos-Flier, J. S. Flier, M. V. Kokoeva, M. Placzek, α-Tanycytes of the adult hypothalamic third ventricle include distinct populations of FGF-responsive neural progenitors. Nat. Commun. 4, 2049 (2013).

5. N. Haan, T. Goodman, A. Najdi-Samiei, C. M. Stratford, R. Rice, E. El Agha, S. Bellusci, M. K. Hajihosseini, Fgf10-expressing tanycytes add new neurons to the appetite/energy-balance regulating centers of the postnatal and adult hypothalamus. J. Neurosci. 33, 6170–6180 (2013).

6. T. Goodman, S. G. Nayar, S. Clare, M. Mikolajczak, R. Rice, S. Mansour, S. Bellusci, M. K. Hajihosseini, Fibroblast growth factor 10 is a negative regulator of postnatal neurogenesis in the mouse hypothalamus. Development. 147 (2020), doi:10.1242/dev.180950.

7. D. A. Lee, S. Yoo, T. Pak, J. Salvatierra, E. Velarde, S. Aja, S. Blackshaw, Dietary and sex-specific factors regulate hypothalamic neurogenesis in young adult mice. Front. Neurosci. 8, 157 (2014).

8. M. Migaud, L. Butrille, M. Batailler, Seasonal regulation of structural plasticity and neurogenesis in the adult mammalian brain: focus on the sheep hypothalamus. Front. Neuroendocrinol. 37, 146–157 (2015).

9. S. Yoo, S. Blackshaw, Regulation and function of neurogenesis in the adult mammalian hypothalamus. Prog. Neurobiol. 170, 53–66 (2018).

10. J. Salvatierra, D. A. Lee, C. Zibetti, M. Duran-Moreno, S. Yoo, E. A. Newman, H. Wang, J. L. Bedont, J. de Melo, A. L. Miranda-Angulo, S. Gil-Perotin, J. M. Garcia-Verdugo, S. Blackshaw, The LIM homeodomain factor Lhx2 is required for hypothalamic tanycyte specification and differentiation. J. Neurosci. 34, 16809–16820 (2014).

11. J. Wan, D. Goldman, Retina regeneration in zebrafish. Curr. Opin. Genet. Dev. 40, 41–47 (2016).

12. M. S. Wilken, T. A. Reh, Retinal regeneration in birds and mice. Curr. Opin. Genet. Dev. 40, 57–64 (2016).

13. A. J. Fischer, R. Bongini, Turning Müller glia into neural progenitors in the retina. Mol. Neurobiol. 42, 199–209 (2010).

14. B. S. Clark, G. L. Stein-O’Brien, F. Shiau, G. H. Cannon, E. Davis-Marcisak, T. Sherman, C. P. Santiago, T. V. Hoang, F. Rajaii, R. E. James-Esposito, R. M. Gronostajski, E. J. Fertig, L. A. Goff, S. Blackshaw, Single-Cell RNA-Seq Analysis of Retinal Development Identifies NFI Factors as Regulating Mitotic Exit and Late-Born Cell Specification. Neuron. 102, 1111–1126.e5 (2019).

15. T. Hoang, J. Wang, P. Boyd, F. Wang, C. Santiago, L. Jiang, S. Yoo, M. Lahne, L. J. Todd, M. Jia, C. Saez, C. Keuthan, I. Palazzo, N. Squires, W. A. Campbell, F. Rajaii, T. Parayil, V. Trinh, D. W. Kim, G. Wang, L. J. Campbell, J. Ash, A. J. Fischer, D. R. Hyde, J. Qian, S. Blackshaw, Gene regulatory networks controlling vertebrate retinal regeneration. Science (2020), doi:10.1126/science.abb8598.

16. D. W. Kim, P. W. Washington, Z. Q. Wang, S. H. Lin, C. Sun, B. T. Ismail, H. Wang, L. Jiang, S. Blackshaw, The cellular and molecular landscape of hypothalamic patterning and differentiation from embryonic to late postnatal development. Nat. Commun. 11, 4360 (2020).

17. R. A. Romanov, E. O. Tretiakov, M. E. Kastriti, M. Zupancic, M. Häring, S. Korchynska, K. Popadin, M. Benevento, P. Rebernik, F. Lallemend, K. Nishimori, F. Clotman, W. D. Andrews, J. G. Parnavelas, M. Farlik, C. Bock, I. Adameyko, T. Hökfelt, E. Keimpema, T. Harkany, Molecular design of hypothalamus development. Nature. 582, 246–252 (2020).

18. T. Pak, S. Yoo, A. L. Miranda-Angulo, H. Wang, S. Blackshaw, Rax-CreERT2 knock-in mice: a tool for selective and conditional gene deletion in progenitor cells and radial glia of the retina and hypothalamus. PLoS One. 9, e90381 (2014).

19. S. Yoo, D. Cha, D. W. Kim, T. V. Hoang, S. Blackshaw, Tanycyte-Independent Control of Hypothalamic Leptin Signaling. Front. Neurosci. 13, 240 (2019).

20. A. Mo, E. A. Mukamel, F. P. Davis, C. Luo, G. L. Henry, S. Picard, M. A. Urich, J. R. Nery, T. J. Sejnowski, R. Lister, S. R. Eddy, J. R. Ecker, J. Nathans, Epigenomic Signatures of Neuronal Diversity in the Mammalian Brain. Neuron. 86, 1369–1384 (2015).

21. R. Chen, X. Wu, L. Jiang, Y. Zhang, Single-Cell RNA-Seq Reveals Hypothalamic Cell Diversity. Cell Rep. 18, 3227–3241 (2017).

22. F. Langlet, Tanycyte Gene Expression Dynamics in the Regulation of Energy Homeostasis. Front. Endocrinol.. 10, 286 (2019).

23. J. N. Campbell, E. Z. Macosko, H. Fenselau, T. H. Pers, A. Lyubetskaya, D. Tenen, M. Goldman, A. M. J. Verstegen, J. M. Resch, S. A. McCarroll, E. D. Rosen, B. B. Lowell, L. T. Tsai, A molecular census of arcuate hypothalamus and median eminence cell types. Nat. Neurosci. 20, 484–496 (2017).

24. E. Matuzelski, J. Bunt, D. Harkins, J. W. C. Lim, R. M. Gronostajski, L. J. Richards, L. Harris, M. Piper, Transcriptional regulation of Nfix by NFIB drives astrocytic maturation within the developing spinal cord. Dev. Biol. 432, 286–297 (2017).

25. S. M. Glasgow, W. Zhu, C. C. Stolt, T.-W. Huang, F. Chen, J. J. LoTurco, J. L. Neul, M. Wegner, C. Mohila, B. Deneen, Mutual antagonism between Sox10 and NFIA regulates diversification of glial lineages and glioma subtypes. Nat. Neurosci. 17, 1322–1329 (2014).

26. D. Vidovic, R. A. Davila, R. M. Gronostajski, T. J. Harvey, M. Piper, Transcriptional regulation of ependymal cell maturation within the postnatal brain. Neural Dev. 13, 2 (2018).

27. B. Kaminskas, T. Goodman, A. Hagan, S. Bellusci, D. M. Ornitz, M. K. Hajihosseini, Characterisation of endogenous players in fibroblast growth factor-regulated functions of hypothalamic tanycytes and energy-balance nuclei. Journal of Neuroendocrinology. 31 (2019),, doi:10.1111/jne.12750.

28. G. La Manno, R. Soldatov, A. Zeisel, E. Braun, H. Hochgerner, V. Petukhov, K. Lidschreiber, M. E. Kastriti, P. Lönnerberg, A. Furlan, J. Fan, L. E. Borm, Z. Liu, D. van Bruggen, J. Guo, X. He, R. Barker, E. Sundström, G. Castelo-Branco, P. Cramer, I. Adameyko, S. Linnarsson, P. V. Kharchenko, RNA velocity of single cells. Nature. 560, 494–498 (2018).

29. G. K. Dhoot, M. K. Gustafsson, X. Ai, W. Sun, D. M. Standiford, C. P. Emerson Jr, Regulation of Wnt signaling and embryo patterning by an extracellular sulfatase. Science. 293, 1663–1666 (2001).

30. D. Mizrak, N. S. Bayin, J. Yuan, Z. Liu, R. M. Suciu, M. J. Niphakis, N. Ngo, K. M. Lum, B. F. Cravatt, A. L. Joyner, P. A. Sims, Single-Cell Profiling and SCOPE-Seq Reveal Lineage Dynamics of Adult Ventricular-Subventricular Zone Neurogenesis and NOTUM as a Key Regulator. Cell Rep. 31, 107805 (2020).

31. R. M. Suciu, A. B. Cognetta 3rd, Z. E. Potter, B. F. Cravatt, Selective Irreversible Inhibitors of the Wnt-Deacylating Enzyme NOTUM Developed by Activity-Based Protein Profiling. ACS Med. Chem. Lett. 9, 563–568 (2018).

32. X. Wang, X.-L. Chou, L. I. Zhang, H. W. Tao, Zona Incerta: An Integrative Node for Global Behavioral Modulation. Trends Neurosci. 43, 82–87 (2020).

33. J. D. Welch, V. Kozareva, A. Ferreira, C. Vanderburg, C. Martin, E. Z. Macosko, Single-Cell Multi-omic Integration Compares and Contrasts Features of Brain Cell Identity. Cell. 177 (2019), pp. 1873–1887.e17.

34. J. Liu, C. Gao, J. Sodicoff, V. Kozareva, E. Z. Macosko, J. D. Welch, Jointly defining cell types from multiple single-cell datasets using LIGER. Nat. Protoc. (2020), doi:10.1038/s41596-020-0391-8.

35. T. Hübschle, E. Thom, A. Watson, J. Roth, S. Klaus, W. Meyerhof, Leptin-induced nuclear translocation of STAT3 immunoreactivity in hypothalamic nuclei involved in body weight regulation. J. Neurosci. 21, 2413–2424 (2001).

36. C. Vaisse, J. L. Halaas, C. M. Horvath, J. E. Darnell Jr, M. Stoffel, J. M. Friedman, Leptin activation of Stat3 in the hypothalamus of wild-type and ob/ob mice but not db/db mice. Nat. Genet. 14, 95–97 (1996).

37. K. Liu, J. Kim, D. W. Kim, Y. S. Zhang, H. Bao, M. Denaxa, S.-A. Lim, E. Kim, C. Liu, I. R. Wickersham, V. Pachnis, S. Hattar, J. Song, S. P. Brown, S. Blackshaw, Lhx6-positive GABA-releasing neurons of the zona incerta promote sleep. Nature. 548, 582–587 (2017).

38. J. A. Dimicco, D. V. Zaretsky, The dorsomedial hypothalamus: a new player in thermoregulation. Am. J. Physiol. Regul. Integr. Comp. Physiol. 292, R47–63 (2007).

39. J. A. DiMicco, B. C. Samuels, M. V. Zaretskaia, D. V. Zaretsky, The dorsomedial hypothalamus and the response to stress: part renaissance, part revolution. Pharmacol. Biochem. Behav. 71, 469–480 (2002).

40. J. Tchieu, E. L. Calder, S. R. Guttikonda, E. M. Gutzwiller, K. A. Aromolaran, J. A. Steinbeck, P. A. Goldstein, L. Studer, NFIA is a gliogenic switch enabling rapid derivation of functional human astrocytes from pluripotent stem cells. Nat. Biotechnol. 37, 267–275 (2019).

41. B. Deneen, R. Ho, A. Lukaszewicz, C. J. Hochstim, R. M. Gronostajski, D. J. Anderson, The transcription factor NFIA controls the onset of gliogenesis in the developing spinal cord. Neuron. 52, 953–968 (2006).

42. Y. Lu, F. Shiau, W. Yi, S. Lu, Q. Wu, J. D. Pearson, A. Kallman, S. Zhong, T. Hoang, Z. Zuo, F. Zhao, M. Zhang, N. Tsai, Y. Zhuo, S. He, J. Zhang, G. L. Stein-O’Brien, T. D. Sherman, X. Duan, E. J. Fertig, L. A. Goff, D. J. Zack, J. T. Handa, T. Xue, R. Bremner, S. Blackshaw, X. Wang, B. S. Clark, Single-Cell Analysis of Human Retina Identifies Evolutionarily Conserved and Species-Specific Mechanisms Controlling Development. Dev. Cell. 53, 473–491.e9 (2020).

43. V. Muthu, H. Eachus, P. Ellis, S. Brown, M. Placzek, Rx3 and Shh direct anisotropic growth and specification in the zebrafish tuberal/anterior hypothalamus. Development. 143, 2651–2663 (2016).

44. T. Shimogori, D. A. Lee, A. Miranda-Angulo, Y. Yang, H. Wang, L. Jiang, A. C. Yoshida, A. Kataoka, H. Mashiko, M. Avetisyan, L. Qi, J. Qian, S. Blackshaw, A genomic atlas of mouse hypothalamic development. Nat. Neurosci. 13, 767–775 (2010).

45. T. S. Corman, S. E. Bergendahl, D. J. Epstein, Distinct temporal requirements for Sonic hedgehog signaling in development of the tuberal hypothalamus. Development. 145 (2018), doi:10.1242/dev.167379.

46. E. A. Newman, D. Wu, M. M. Taketo, J. Zhang, S. Blackshaw, Canonical Wnt signaling regulates patterning, differentiation and nucleogenesis in mouse hypothalamus and prethalamus. Dev. Biol. 442, 236–248 (2018).

47. X. Wang, D. Kopinke, J. Lin, A. D. McPherson, R. N. Duncan, H. Otsuna, E. Moro, K. Hoshijima, D. J. Grunwald, F. Argenton, C.-B. Chien, L. C. Murtaugh, R. I. Dorsky, Wnt signaling regulates postembryonic hypothalamic progenitor differentiation. Dev. Cell. 23, 624–636 (2012).

48. J. E. Lee, S.-F. Wu, L. M. Goering, R. I. Dorsky, Canonical Wnt signaling through Lef1 is required for hypothalamic neurogenesis. Development. 133, 4451–4461 (2006).

49. A. P. Jadhav, S.-H. Cho, C. L. Cepko, Notch activity permits retinal cells to progress through multiple progenitor states and acquire a stem cell property. Proceedings of the National Academy of Sciences. 103 (2006), pp. 18998–19003.

50. J. P. Magnusson, C. Göritz, J. Tatarishvili, D. O. Dias, E. M. K. Smith, O. Lindvall, Z. Kokaia, J. Frisén, A latent neurogenic program in astrocytes regulated by Notch signaling in the mouse. Science. 346, 237–241 (2014).

51. L. Todd, I. Palazzo, N. Squires, N. Mendonca, A. J. Fischer, BMP-and TGFβ-signaling regulate the formation of Müller glia-derived progenitor cells in the avian retina. Glia. 65, 1640–1655 (2017).

52. J. Stipursky, D. Francis, R. S. Dezonne, A. P. Bérgamo de Araújo, L. Souza, C. A. Moraes, F. C. Alcantara Gomes, TGF-β1 promotes cerebral cortex radial glia-astrocyte differentiation in vivo. Front. Cell. Neurosci. 8, 393 (2014).

53. R. A. Poché, Y. Furuta, M.-C. Chaboissier, A. Schedl, R. R. Behringer, Sox9 is expressed in mouse multipotent retinal progenitor cells and functions in Müller glial cell development. J. Comp. Neurol. 510, 237–250 (2008).

54. C. C. Stolt, P. Lommes, E. Sock, M.-C. Chaboissier, A. Schedl, M. Wegner, The Sox9 transcription factor determines glial fate choice in the developing spinal cord. Genes Dev. 17, 1677–1689 (2003).

55. S. Aslanpour, J. M. Rosin, A. Balakrishnan, N. Klenin, F. Blot, G. Gradwohl, C. Schuurmans, D. M. Kurrasch, Ascl1 is required to specify a subset of ventromedial hypothalamic neurons. Development. 147 (2020), p. dev180067.

56. N. L. Jorstad, M. S. Wilken, L. Todd, C. Finkbeiner, P. Nakamura, N. Radulovich, M. J. Hooper, A. Chitsazan, B. A. Wilkerson, F. Rieke, T. A. Reh, STAT Signaling Modifies Ascl1 Chromatin Binding and Limits Neural Regeneration from Muller Glia in Adult Mouse Retina. Cell Rep. 30, 2195–2208.e5 (2020).

57. N. L. Jorstad, M. S. Wilken, W. N. Grimes, S. G. Wohl, L. S. VandenBosch, T. Yoshimatsu, R. O. Wong, F. Rieke, T. A. Reh, Stimulation of functional neuronal regeneration from Müller glia in adult mice. Nature. 548, 103–107 (2017).

58. W. Wang, D. Mullikin-Kilpatrick, J. E. Crandall, R. M. Gronostajski, E. D. Litwack, D. L. Kilpatrick, Nuclear factor I coordinates multiple phases of cerebellar granule cell development via regulation of cell adhesion molecules. J. Neurosci. 27, 6115–6127 (2007).

59. M. Piper, L. Harris, G. Barry, Y. H. E. Heng, C. Plachez, R. M. Gronostajski, L. J. Richards, Nuclear factor one X regulates the development of multiple cellular populations in the postnatal cerebellum. J. Comp. Neurol. 519, 3532–3548 (2011).

60. J. R. Ryu, C. J. Hong, J. Y. Kim, E.-K. Kim, W. Sun, S.-W. Yu, Control of adult neurogenesis by programmed cell death in the mammalian brain. Molecular Brain. 9 (2016),, doi:10.1186/s13041-016-0224-4.

61. A. Lafzi, C. Moutinho, S. Picelli, H. Heyn, Tutorial: guidelines for the experimental design of single-cell RNA sequencing studies. Nat. Protoc. 13, 2742–2757 (2018).

62. P. Vercruysse, D. Vieau, D. Blum, Å. Petersén, L. Dupuis, Hypothalamic Alterations in Neurodegenerative Diseases and Their Relation to Abnormal Energy Metabolism. Front. Mol. Neurosci. 11, 2 (2018).

63. J. P. Thaler, C.-X. Yi, E. A. Schur, S. J. Guyenet, B. H. Hwang, M. O. Dietrich, X. Zhao, D. A. Sarruf, V. Izgur, K. R. Maravilla, H. T. Nguyen, J. D. Fischer, M. E. Matsen, B. E. Wisse, G. J. Morton, T. L. Horvath, D. G. Baskin, M. H. Tschöp, M. W. Schwartz, Obesity is associated with hypothalamic injury in rodents and humans. J. Clin. Invest. 122, 153–162 (2012).

64. Y.-C. Hsu, J. Osinski, C. E. Campbell, E. D. Litwack, D. Wang, S. Liu, C. J. Bachurski, R. M. Gronostajski, Mesenchymal nuclear factor I B regulates cell proliferation and epithelial differentiation during lung maturation. Dev. Biol. 354, 242–252 (2011).

65. C. E. Campbell, M. Piper, C. Plachez, Y.-T. Yeh, J. S. Baizer, J. M. Osinski, E. D. Litwack, L. J. Richards, R. M. Gronostajski, The transcription factor Nfix is essential for normal brain development. BMC Dev. Biol. 8, 52 (2008).

66. C. Zeng, F. Pan, L. A. Jones, M. M. Lim, E. A. Griffin, Y. I. Sheline, M. A. Mintun, D. M. Holtzman, R. H. Mach, Evaluation of 5-ethynyl-2’-deoxyuridine staining as a sensitive and reliable method for studying cell proliferation in the adult nervous system. Brain Res. 1319, 21–32 (2010).

67. N. Pentinmikko, S. Iqbal, M. Mana, S. Andersson, A. B. Cognetta 3rd, R. M. Suciu, J. Roper, K. Luopajärvi, E. Markelin, S. Gopalakrishnan, O.-P. Smolander, S. Naranjo, T. Saarinen, A. Juuti, K. Pietiläinen, P. Auvinen, A. Ristimäki, N. Gupta, T. Tammela, T. Jacks, D. M. Sabatini, B. F. Cravatt, Ö. H. Yilmaz, P. Katajisto, Notum produced by Paneth cells attenuates regeneration of aged intestinal epithelium. Nature. 571, 398–402 (2019).

68. S. Yoo, D. Cha, S. Kim, L. Jiang, P. Cooke, M. Adebesin, A. Wolfe, R. Riddle, S. Aja, S. Blackshaw, Tanycyte ablation in the arcuate nucleus and median eminence increases obesity susceptibility by increasing body fat content in male mice. Glia. 68, 1987–2000 (2020).

69. M. Schneeberger, L. Parolari, T. Das Banerjee, V. Bhave, P. Wang, B. Patel, T. Topilko, Z. Wu, C. H. J. Choi, X. Yu, K. Pellegrino, E. A. Engel, P. Cohen, N. Renier, J. M. Friedman, A. R. Nectow, Regulation of Energy Expenditure by Brainstem GABA Neurons. Cell. 178, 672–685.e12 (2019).

70. G. X. Y. Zheng, J. M. Terry, P. Belgrader, P. Ryvkin, Z. W. Bent, R. Wilson, S. B. Ziraldo, T. D. Wheeler, G. P. McDermott, J. Zhu, M. T. Gregory, J. Shuga, L. Montesclaros, J. G. Underwood, D. A. Masquelier, S. Y. Nishimura, M. Schnall-Levin, P. W. Wyatt, C. M. Hindson, R. Bharadwaj, A. Wong, K. D. Ness, L. W. Beppu, H. J. Deeg, C. McFarland, K. R. Loeb, W. J. Valente, N. G. Ericson, E. A. Stevens, J. P. Radich, T. S. Mikkelsen, B. J. Hindson, J. H. Bielas, Massively parallel digital transcriptional profiling of single cells. Nat. Commun. 8, 14049 (2017).

71. T. Stuart, A. Butler, P. Hoffman, C. Hafemeister, E. Papalexi, W. M. Mauck, Y. Hao, M. Stoeckius, P. Smibert, R. Satija, Comprehensive Integration of Single-Cell Data. Cell. 177 (2019), pp. 1888–1902.e21.

72. S. L. Wolock, R. Lopez, A. M. Klein, Scrublet: computational identification of cell doublets in single-cell transcriptomic data. Cell Systems. 8, 231–291 (2019).

73. I. Korsunsky, N. Millard, J. Fan, K. Slowikowski, F. Zhang, K. Wei, Y. Baglaenko, M. Brenner, P.-R. Loh, S. Raychaudhuri, Fast, sensitive and accurate integration of single-cell data with Harmony. Nat. Methods. 16, 1289–1296 (2019).

74. V. Bergen, M. Lange, S. Peidli, F. A. Wolf, F. J. Theis, Generalizing RNA velocity to transient cell states through dynamical modeling. Nat. Biotechnol. (2020), doi:10.1038/s41587-020-0591-3.

75. I. Tirosh, B. Izar, S. M. Prakadan, M. H. Wadsworth 2nd, D. Treacy, J. J. Trombetta, A. Rotem, C. Rodman, C. Lian, G. Murphy, M. Fallahi-Sichani, K. Dutton-Regester, J.-R. Lin, O. Cohen, P. Shah, D. Lu, A. S. Genshaft, T. K. Hughes, C. G. K. Ziegler, S. W. Kazer, A. Gaillard, K. E. Kolb, A.-C. Villani, C. M. Johannessen, A. Y. Andreev, E. M. Van Allen, M. Bertagnolli, P. K. Sorger, R. J. Sullivan, K. T. Flaherty, D. T. Frederick, J. Jané-Valbuena, C. H. Yoon, O. Rozenblatt-Rosen, A. K. Shalek, A. Regev, L. A. Garraway, Dissecting the multicellular ecosystem of metastatic melanoma by single-cell RNA-seq. Science. 352, 189–196 (2016).

76. K. Street, D. Risso, R. B. Fletcher, D. Das, J. Ngai, N. Yosef, E. Purdom, S. Dudoit, Slingshot: cell lineage and pseudotime inference for single-cell transcriptomics. BMC Genomics. 19, 477 (2018).

77. X. Qiu, A. Hill, J. Packer, D. Lin, Y.-A. Ma, C. Trapnell, Single-cell mRNA quantification and differential analysis with Census. Nat. Methods. 14, 309–315 (2017).

78. Y. Zhang, T. Liu, C. A. Meyer, J. Eeckhoute, D. S. Johnson, B. E. Bernstein, C. Nusbaum, R. M. Myers, M. Brown, W. Li, X. S. Liu, Model-based analysis of ChIP-Seq (MACS). Genome Biol. 9, R137 (2008).

79. A. N. Schep, B. Wu, J. D. Buenrostro, W. J. Greenleaf, chromVAR: inferring transcription-factor-associated accessibility from single-cell epigenomic data. Nat. Methods. 14, 975–978 (2017).

80. M. R. Corces, J. M. Granja, S. Shams, B. H. Louie, J. A. Seoane, W. Zhou, T. C. Silva, C. Groeneveld, C. K. Wong, S. W. Cho, A. T. Satpathy, M. R. Mumbach, K. A. Hoadley, A. G. Robertson, N. C. Sheffield, I. Felau, M. A. A. Castro, B. P. Berman, L. M. Staudt, J. C. Zenklusen, P. W. Laird, C. Curtis, Cancer Genome Atlas Analysis Network, W. J. Greenleaf, H. Y. Chang, The chromatin accessibility landscape of primary human cancers. Science. 362 (2018), doi:10.1126/science.aav1898.

81. Z. Shao, Y. Zhang, G.-C. Yuan, S. H. Orkin, D. J. Waxman, MAnorm: a robust model for quantitative comparison of ChIP-Seq data sets. Genome Biol. 13, R16 (2012).

82. S. Heinz, C. Benner, N. Spann, E. Bertolino, Y. C. Lin, P. Laslo, J. X. Cheng, C. Murre, H. Singh, C. K. Glass, Simple Combinations of Lineage-Determining Transcription Factors Prime cis-Regulatory Elements Required for Macrophage and B Cell Identities. Molecular Cell. 38 (2010), pp. 576–589.

83. J. Wang, C. Zibetti, P. Shang, S. R. Sripathi, P. Zhang, M. Cano, T. Hoang, S. Xia, H. Ji, S. L. Merbs, D. J. Zack, J. T. Handa, D. Sinha, S. Blackshaw, J. Qian, ATAC-Seq analysis reveals a widespread decrease of chromatin accessibility in age-related macular degeneration. Nat. Commun. 9, 1364 (2018).

84. H. A. Pliner, J. S. Packer, J. L. McFaline-Figueroa, D. A. Cusanovich, R. M. Daza, D. Aghamirzaie, S. Srivatsan, X. Qiu, D. Jackson, A. Minkina, A. C. Adey, F. J. Steemers, J. Shendure, C. Trapnell, Cicero Predicts cis-Regulatory DNA Interactions from Single-Cell Chromatin Accessibility Data. Mol. Cell. 71, 858–871.e8 (2018).

85. E. Eden, R. Navon, I. Steinfeld, D. Lipson, Z. Yakhini, GOrilla: a tool for discovery and visualization of enriched GO terms in ranked gene lists. BMC Bioinformatics. 10, 48 (2009).

86. F. Supek, M. Bošnjak, N. Škunca, T. Šmuc, REVIGO summarizes and visualizes long lists of gene ontology terms. PLoS One. 6, e21800 (2011).

87. J. E. Frandolig, C. J. Matney, K. Lee, J. Kim, M. Chevée, S.-J. Kim, A. A. Bickert, S. P. Brown, The Synaptic Organization of Layer 6 Circuits Reveals Inhibition as a Major Output of a Neocortical Sublamina. Cell Rep. 28, 3131–3143.e5 (2019).

